# Batch Effects Remain a Fundamental Barrier to Universal Embeddings in Single-Cell Foundation Models

**DOI:** 10.64898/2025.12.19.695371

**Authors:** Linting Wang, Chihao Zhang, Shihua Zhang

## Abstract

Constructing a cell universe requires integrating heterogeneous single-cell RNA-seq datasets, but is hindered by diverse batch effects. Single-cell foundation models (scFMs), inspired by large language models, aim to learn universal cellular embeddings from large-scale single-cell data. However, unlike language, single-cell data are sparse, noisy, and strongly affected by batch effects that limit cross-dataset transferability. Our systematic evaluation across diverse batch scenarios reveals that current scFMs fail to effectively remove batch effects, with batch signals persisting in pretrained embeddings. Post-hoc batch-centering partially improves alignment, highlighting the need for future scFMs to integrate explicit batch-effect correction mechanisms to achieve true universal cellular embeddings.

## Introduction

Biological systems are complex yet functionally unified entities. In the human body, organs such as the brain, heart, lungs, and liver maintain precise coordination through cross-organ cellular communication networks, including cellular functions, intercellular signaling, and dynamic cellular states [1-3].

Therefore, achieving a holistic understanding of biological systems requires analyses that extend beyond individual organs and create a “cell universe” (**Fig. 1a**). We define this as a unified embedding space where cells are organized solely by their biological identity: shared cell types from different organs cluster together, whereas organ-specific cell populations preserve their distinct biological signature. This space could depict cellular diversity and facilitate systematic analyses of development, communication, and disease progression across the entire organism.

**Figure 1.**
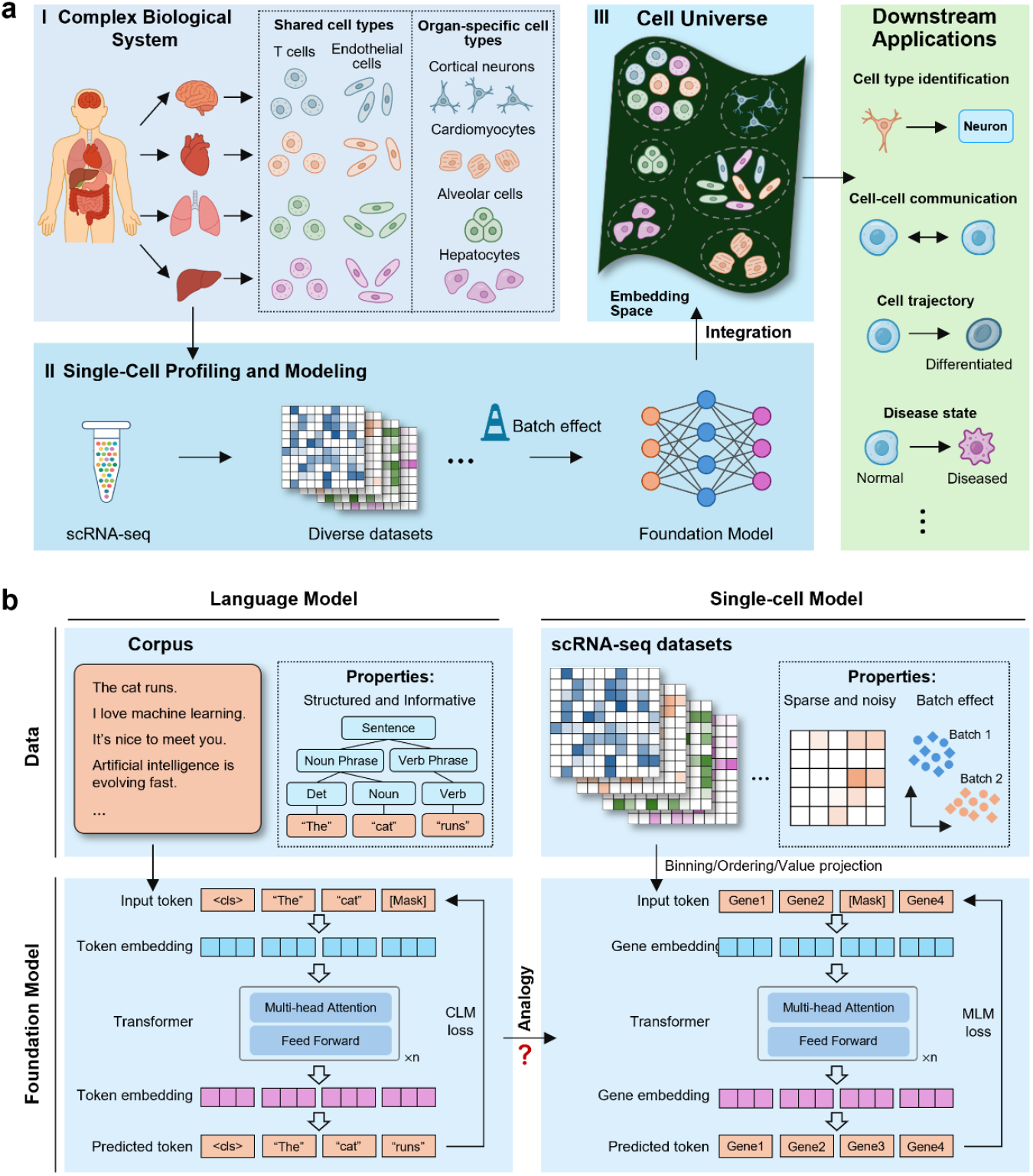
Conceptual overview of constructing the cell universe and comparison between language and single-cell foundational models (scFMs). **a**, (I) Complex biological systems consist of multiple organs containing both shared and organ-specific cell types. (II) Single-cell profiling and modeling capture these diverse cellular states using scRNA-seq, followed by data integration to build a cell universe. (III) Shared and organ-specific cells are represented in an embedding space, where shared cells cluster together and organ-specific cells separate. (IV) This space enables multiple downstream analyses, such as cell type identification. **b**, For language, structured and informative text corpora are tokenized and encoded into embeddings for next-token prediction. In contrast, single-cell data are sparse, noisy, and affected by batch effects. Genes are treated as tokens with expression and position embeddings, and models are trained via masked gene modeling to learn biological embeddings that are analogous to those in language models. MLM, masked language modeling; CLM, causal language modeling.

Single-cell RNA sequencing (scRNA-seq) has made this vision attainable by profiling gene expression at single-cell resolution [4-6]. Yet, individual experiments capture only limited subsets of cells under distinct technical and biological conditions, resulting in fragmented data landscapes. To overcome this, large international initiatives, such as the Human Cell Atlas, aim to integrate diverse datasets into a comprehensive reference map [7]. However, batch effects, the systematic variation across experiments [8-11], remain a critical barrier, preventing cells from being compared within a shared representation space.

Large language models (LLMs) in natural language processing (NLP) have demonstrated the ability to learn a linguistic embedding space where words, phrases, or sentences with similar meanings cluster together [12-14]. Inspired by this success, single-cell foundation models (scFMs) have emerged as a promising paradigm for creating a “cell universe”. These scFMs, such as scBERT, Geneformer, scGPT, scFoundation, and Nicheformer [15-27], typically adopt the Transformer architecture and are pretrained on large-scale datasets to learn universal cell embeddings (**Fig. 1b**).

However, single-cell data differ fundamentally from language data (**Table 1** and **Fig. 1b**). While language is structured, dense, and semantically coherent, single-cell data are high-dimensional, sparse, and confounded by substantial batch effects. Consequently, Transformers designed for text may struggle to capture the intrinsic properties of single-cell data, limiting the practical utility of current scFMs. Recent benchmarks reveal that the current scFMs [28-33] often underperform traditional computational methods or even simple baselines in zero-shot batch correction [28, 32, 33] and perturbation prediction [29, 31]. These findings suggest that current scFMs still fail to learn universal cell embeddings across heterogeneous datasets. To improve scFMs, recent reviews suggest strategies such as data expansion or objective refinement [28, 34-39]. However, these studies primarily treat batch correction as a downstream task, rather than recognizing batch-invariance as a fundamental prerequisite for scFMs.

**Table 1.** Comparative characteristics of language data and single-cell data. MLM, masked language modeling; CLM, causal language modeling; NLP, natural language processing.

|  | Language data | Single cell data |
| --- | --- | --- |
| Data type | Sequential data | Tabular data |
| Knowledge level | Compressed knowledge | Raw data |
| Noise level | Low | High |
| Data structure | Highly structured with strong contextual dependencies | Highly sparse with low information density |
| Batch effect | Low | High |
| Data scale | 100B~10TB tokens | 10M~100M cells |
| Model size | 10B~1TB parameters | 10M~1B parameters |
| Token definition | Words, subwords, or characters | Genes or discretized expression |
| Input strategy | Embedded after tokenization | Binning, ordering or value projection |
| Loss function | CLM or MLM | Mainly MLM |
| Downstream applications | Broad NLP tasks (e.g., summarization, question answering, translation) | Biological tasks such as cell type annotation and perturbation prediction |

In contrast, we argue that a practical scFM must generate batch-invariant cell embeddings, which are necessary to provide universal cellular embeddings and create a veritable cell universe. In this study, we systematically evaluated existing scFMs on their ability to remove batch effects across diverse sources of variation. We found that scFMs fail to eliminate batch differences, with substantial batch-specific information remaining in cell embeddings. That contrasts sharply with LLMs, whose latent spaces form stable semantic structures largely independent of language identity. Although a simple batch-centering step partially improved alignment, it remained insufficient. Together, these results suggest that current scFMs have not yet achieved effective batch effect removal. We propose that future scFMs should move beyond direct analogies to language models and incorporate explicit mechanisms to correct batch effects.

## Results

### Cell Embeddings from scFMs Exhibit Pronounced Batch Effects

We first evaluated the ability of existing scFMs to remove batch effects. Building on prior evaluations [28, 32, 33], we extended the benchmarking framework to cover multiple sources of batch variation, including tissue origin, inter-individual differences, disease state, assay type, RNA source, and species, which represent the diverse complexity of batch effects. Evaluation datasets were obtained from the CELLxGENE database [40] and summarized in **Supplementary Table S1**. The recently released immune and liver datasets were not included in the pretraining data for any scFMs, thereby minimizing potential information leakage. For each dataset, we extracted zero-shot cell embeddings from eight scFMs, including Geneformer [16, 17], scGPT [18], scFoundation [19], GeneCompass [21], CellPLM [22], UCE [23], Nicheformer [24], and scCello [26]. We compared them with traditional batch integration methods, including Harmony [45], scVI [46], and Scanorama [47]. We included PCA as a linear baseline.

The UMAP visualizations (**Fig. 2a,Supplementary Figs. S1-S6**) showed that embeddings generated by scFMs exhibited pronounced batch effects across all scenarios. These observations were quantitatively supported by the aggregated scIB score [8], a metric designed to evaluate batch mixing and cell-type conservation in single-cell embeddings jointly (**Fig. 2b**). Except for scCello, scFMs consistently underperformed Harmony and scVI, and in some cases even fell below the PCA baseline. Notably, for datasets with mild batch effects, such as tissues and diseases, scFMs performed comparably to traditional methods. But, as batch heterogeneity increased, particularly in assay and RNA source scenarios, the performance gap widened substantially. In summary, these findings are consistent with prior studies [28, 32, 36] and show that current scFMs have not yet effectively eliminated batch effects, particularly in datasets with severe batch variation.

**Figure 2.**
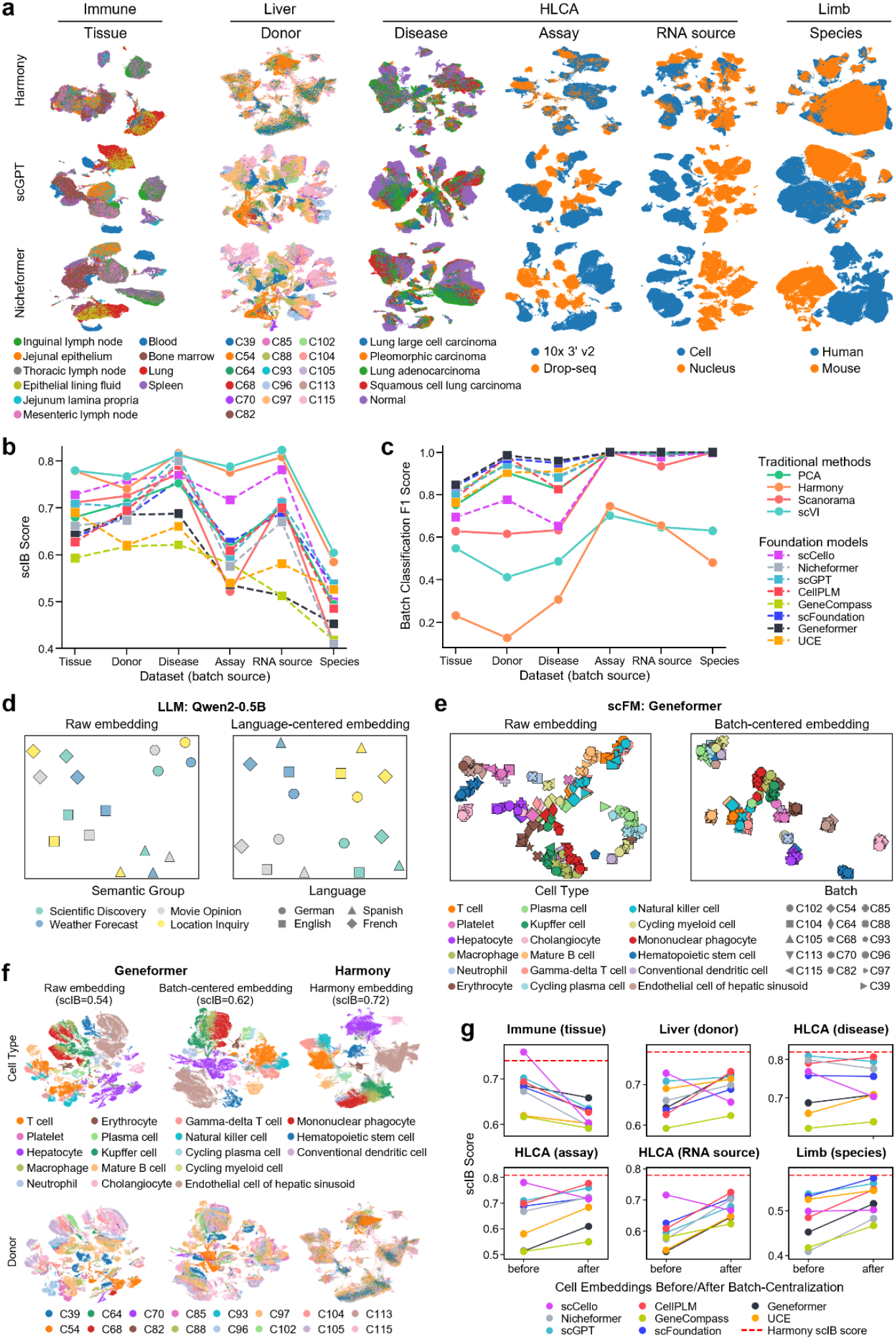
Cross-dataset evaluation of batch effects in scFMs. **a**, UMAP visualizations of six benchmark datasets using cell embeddings derived from the traditional integration method Harmony and two representative scFMs (scGPT and Nicheformer). See results for other models in Fig. S1–S6. **b-c**, Quantitative differences of traditional approaches and scFMs based on (b) scIB scores and (c) batch-classification F1 scores across six datasets that represent distinct batch sources. **d**, UMAP visualizations of raw and language-centered sentence embeddings generated by Qwen2-0.5B on a multilingual corpus, illustrating semantic alignment through language centralization. **e**, UMAP visualizations of raw and batch-centered embeddings generated by Geneformer on the Liver dataset. **f**, Differences of raw and batch-centered Geneformer embeddings with Harmony embeddings on the Liver dataset, visualized by cell type (top) and donor (bottom). **g**, Changes of scIB scores before and after batch-centralization across six datasets for eight scFMs.

Next, we examined the types of information retained within learned embeddings of scFMs. We performed a probing analysis, a widely used approach in machine learning interpretability studies [48, 49]. We trained two logistic regression probes on the embeddings from each method to predict cell type and batch separately, using five-fold cross-validation. High predictive accuracy indicates that the embeddings explicitly encode the corresponding information. As shown in **Fig. 2c** and **Supplementary Fig. S7**, although all embeddings achieved high cell-type prediction accuracy and macro-F1 score, those generated by scFMs consistently exhibited higher batch-prediction accuracy than traditional integration methods and the PCA baseline. These findings further demonstrate that the embeddings generated by current scFMs still retain substantial batch-related information.

### Comparison with language models highlights the limitations of scFM

The design of scFMs is heavily inspired by the Transformers in NLP, with most models analogizing “genes” to “words” and “cells” to “sentences” [15, 16, 18, 38]. This raises a natural question: do language models exhibit a phenomenon analogous to the “batch effect” observed in single-cell data?

To investigate this, we first examined the structure of the sentence embedding space in the open-source LLM Qwen [50, 51]. We extracted embeddings for semantically equivalent sentences across four languages and visualized them using UMAP (**Fig. 2d,left** and **Supplementary Fig. S8**). The results show that language identity dominates in embedding. Sentences of the same language are closely clustered, while sentences of different languages are significantly separated. Inspired by prior work [52, 53], we then subtracted a “language embedding” from each sentence and found that semantically equivalent sentences clustered together (**Fig. 2d,right** and **Supplementary Fig. S8**). This discovery suggests that LLMs indeed encode shared semantic embeddings, which were initially masked by signals of specific languages.

We next investigated whether scFMs exhibit a similar embedding structure.

For each dataset, we computed the average embedding for each cell type within batches and found that these “cell-type embeddings” were strongly affected by batch effects (**Fig. 2e**, left). We then subtracted the batch-specific mean embedding from the embeddings of each cell type. However, unlike in LLMs, this adjustment only partially mitigated batch effects and failed to achieve global alignment of the same cell type across batches.

In summary, these findings reveal a fundamental distinction: while LLMs can recover stable semantic structures once language identity is removed, scFMs have not yet uncovered biologically universal cell embeddings. This shows the limitations of the “language analogy” in modeling single-cell data and underscores the need to incorporate explicit batch-correction mechanisms.

### Batch-centralization partially reduced the batch effect of scFM embeddings

To illustrate the importance of batch-effect removal in scFMs design, we applied a simple batch-centralization adjustment by subtracting the mean embedding of each batch from individual cells. This adjustment visibly reduced batch effects in UMAP projections (**Fig. 2f**; **Supplementary Figs. S9–S14**) and improved scIB scores for most models and datasets (**Fig. 2g**). Probing analyses showed decreased batch-prediction accuracy with largely preserved cell-type prediction (**Supplementary Fig. S15**).

However, the effect of this linear adjustment was limited. The scIB score decreased for scCello, as well as in datasets of tissue and disease batch scenarios (**Fig. 2g**), indicating potential overcorrection. In addition, the batch-centered embeddings of scFMs still failed to outperform traditional methods (**Fig. 2g** and **Supplementary Figs. S9-S14**). These results show that while the simple post-hoc linear adjustment can partially mitigate batch effects, it remains insufficient, highlighting the need for architectures and training objectives that explicitly address batch variation.

## Discussion

Foundation models hold great promise for constructing a cell universe. However, our evaluation indicates that current scFMs still exhibit substantial batch effects, limiting their robustness and generalization. The central goal of an scFM is to learn universal cell embeddings that capture fundamental biological knowledge while remaining consistent across experiments, tissues, and even species [23]. Such a model should embed heterogeneous single-cell data into a shared latent space without batch effects. Therefore, removing batch effects is a fundamental prerequisite for achieving true universal cell embeddings.

Notably, most existing scFMs lack explicit batch correction mechanisms (**Table S2**). Only a few models incorporate related designs. For example, CellPLM [22] introduces batch embeddings in its decoder; scCello [26] and NephrobaseCell+ [27] leverage supervised learning with cell types; NephrobaseCell+ further employs adversarial learning to suppress batch-related signals; Nicheformer [24] incorporates additional modality and assay tokens. Despite these efforts, CellPLM and Nicheformer still underperformed Harmony and scVI (**Fig. 2b**), indicating that such strategies are insufficient. scCello performed best among scFMs, demonstrating the benefit of supervised learning, but it still lags behind traditional algorithms in batch mixing (**Supplementary Figs. S1-S6**). Therefore, most scFMs still rely on implicit alignment across batches during pretraining.

However, this implicit alignment is fundamentally limited due to structural differences between language and single-cell data (**Table 1**). The language is dense, low-noise, and governed by syntactic rules, whereas scRNA-seq data are high-dimensional, sparse, noisy, and highly heterogeneous across batches [8-11]. This mismatch also reflects a deeper semantic gap: language represents compressed human knowledge, whereas single-cell profiles remain at the data level, lacking the structural biological prior. As a result, Transformer architectures, which were originally designed for textual semantics, are not naturally aligned with the characteristics of single-cell data. The high dimensionality and sparsity of single-cell profiles necessitate substantial preprocessing before model input, leading to information loss [30]. The noise in single-cell data makes it difficult for the model to capture biological signals. Moreover, common self-supervised objectives (e.g., masked token recovery) tend to capture batch-specific signals instead of removing them. Therefore, directly applying Transformer-based architectures to single-cell data faces challenges in capturing their intrinsic features.

Although LLMs benefit from scaling laws, we found no analogous relationship between model size or training data volume and batch-effect removal performance across current scFMs (**Supplementary Fig. S16**), suggesting that scale is not the primary limitation for scFMs. Instead, learning strategy appears more critical: scCello, despite its small model size and limited training data, achieved better batch correction performance than other scFMs. Its supervised design directly targets biologically meaningful variation, producing embeddings that are more robust to technical noise.

Therefore, future scFMs should explicitly address batch effects in their architecture and training objectives. We propose several strategies to achieve this: (1) **Input-level design:** introduce explicit batch tokens, technical metadata, or contextual encodings. Our results suggest that a separable batch component exists, and this design enables us to identify and remove it. (2) **Objective-level design:** incorporate adversarial objectives, distribution alignment, or invariance constraints into the self-supervised loss to reduce sensitivity to non-biological variation [27, 54]. This is beneficial as our results show that simple linear correction is insufficient. (3) **Supervision-level design:** leverage biological priors such as cell type and regulatory networks as supervision signals [26, 27, 55, 56]. Such priors help prevent overcorrection and preserve biologically meaningful variation.

In summary, addressing batch effects is not merely a technical challenge but a conceptual cornerstone for achieving unified single-cell modeling and constructing a universal cell atlas. Achieving structural separation between biological and technical variation is a prerequisite for genuine single-cell intelligence.

## Methods

### Datasets collection and preprocessing

A summary of all evaluation datasets and their key characteristics is provided in **Supplementary Table S1**.

#### Immune dataset

The human immune atlas [41] comprises over 1.25 million immune cells isolated from 10 tissues, including blood, lymphoid organs, and mucosal tissues of 24 human organ donors. We restricted the analysis to one donor, D529, and used its cells from 10 tissues (85,588 cells in total), ensuring that batch variation primarily arises from tissue differences rather than donor heterogeneity. We set the tissue of origin as the batch key.

#### Liver dataset

The healthy human liver datasets [42] comprise 42,660 cells from 9 pediatric donors and 26,372 cells from 7 adult donors. Donor identity was used as the batch variable to evaluate batch-associated variation across individuals.

#### HLCA dataset

The Human Lung Cell Atlas (HLCA) [43] is a large-scale single-cell reference atlas of the human respiratory system, comprising over 2 million cells from 486 individuals across 49 datasets. To systematically examine different sources of batch variation, we constructed evaluation subsets along three axes:

##### Assay

two datasets using distinct profiling technologies — *Meyer_2019* (10x 3’ v2) and *Schiller_2020* (Drop-seq).

##### RNA source

two datasets differing in RNA origin — *Sun_2020* (snRNA-seq) and *Meyer_2019* (scRNA-seq).

##### Disease state

two datasets representing healthy and pathological conditions — *Thienpont_2018* (four lung cancer types) and *Meyer_2019* (healthy donors).

These subsets enable the assessment of assay-, RNA-source-, and disease-driven batch effects.

#### Limb dataset

The embryonic limb dataset [44] comprises a cross-species single-cell transcriptomic atlas capturing limb development in human and mouse embryos. We mapped the mouse genes to their human orthologs before integration, and employed the species label as the batch key.

### Data preprocessing

All datasets were obtained from the CELLxGENE data portal [40]. Depending on the input requirements of each scFM, we used either the raw count matrix or its normalized counterpart. We normalized the raw counts using ‘sc.pp.normalize_total’ (target_sum=1e4) followed by log-transformation with ‘sc.pp.log1p’ functions from the Scanpy Python package. We removed the cells of unknown cell types. Additional preprocessing required by specific models is detailed in the following sections.

### Traditional integration methods

#### PCA

Principal Component Analysis (PCA) was used as an uncorrected baseline to provide a reference representation of the data without explicit batch-effect adjustment. PCA performs linear dimensionality reduction by projecting high-dimensional gene expression profiles onto orthogonal components that maximize variance. In our analyses, PCA was applied directly to the normalized expression matrix. We selected the highly variable genes using the ‘scib.pp.reduce_data’ function from the scib Python package, and employed the resulting PCA embedding and variable gene set in subsequent integration methods.

#### Harmony

Harmony [45] integrates multiple single-cell datasets by refining an initial PCA embedding through iterative soft clustering, aligning cells across batches while preserving biological structure. We applied Harmony using the ‘scib.integration.harmony’ function with batch labels provided through the designated batch key. We set the other parameters to the defaults. The embedding was initialized with the PCA representation computed previously, using the same highly variable gene set.

#### Scanorama

Scanorama [47] integrates heterogeneous single-cell transcriptomic datasets by identifying mutual nearest neighbors across datasets and performing manifold alignment to generate corrected embeddings. In this study, we applied Scanorama through the ‘scib.integration.scanorama’ function from the scib Python package. We specified the batch key corresponding to each dataset in advance and used the same set of highly variable genes identified using PCA. We set other parameters to the defaults.

#### scVI

scVI [46] is a deep generative variational autoencoder that models gene expression counts probabilistically, incorporates batch covariates, and learns a latent embedding that mitigates technical variation. In this study, we applied the ‘scib.integration.scvi’ function from the scib Python package to integrate datasets, specifying the batch key and using the previously defined highly variable genes. We set other parameters to the defaults.

### Single-cell foundation models (scFMs)

The basic information of current scFMs is summarized in **Supplementary Table S2**.

#### Geneformer

Geneformer [16, 17] employs a transformer-based architecture in which each cell’s transcriptome is converted into a rank-value representation, followed by self-attention layers to capture contextual gene– gene relationships. The V2 version of the model was pretrained on a corpus of approximately 104 million human single-cell profiles. In this study, we used the Geneformer-V2-316M model. We processed the evaluation datasets and tokenized them using ‘tk.tokenize_data’, and obtained the cell embeddings through the ‘EmbExtractor.extract_embs’ function with default parameters.

#### scGPT

scGPT [18] is a generative pre-trained transformer model that treats genes as tokens and each cell as an analog of a sentence. It was pretrained on more than 33 million single-cell profiles collected from diverse healthy human tissues. Gene expression values are discretized via binning before being passed to the model. We adopted the scGPT-human model with default settings to extract embeddings through the ‘scg.tasks.embed_data’ function with default parameters.

#### CellPLM

CellPLM [22] introduces a cell-language model paradigm that explicitly encodes both gene–gene and cell–cell relationships. Its architecture includes a gene-expression embedder, a transformer encoder, a Gaussian-mixture latent space prior, and a batch-aware decoder. We extracted the cell embeddings using the ‘CellEmbeddingPipeline.predict’ function with default parameters.

#### UCE

The Universal Cell Embeddings (UCE) model [23] is a self-supervised foundation model designed to produce universal embeddings across tissues and species, without requiring dataset-specific retraining or gene selection. We generated the cell embeddings for each evaluation dataset using the script ‘eval_single_anndata.py’ with default parameters.

#### scFoundation

scFoundation [19] is a large-scale pretrained model with an asymmetric transformer-like encoder–decoder architecture containing approximately 100 million parameters. It was pretrained on over 50 million human single-cell profiles. It adopts a value-projection strategy to convert continuous expression levels into token embeddings and employs read-depth-aware pretraining tasks to accommodate variation in sequencing depth. We extracted the cell embeddings using the script ‘get_embedding.py’ with default parameters.

#### Nicheformer

Nicheformer [24] uses a transformer-based design to jointly model dissociated single-cell and spatial transcriptomic data across human and mouse. Pretraining was performed on approximately 110 million cells. The model includes additional modality, organism, and assay tokens to condition the embedding space. We generated the cell embeddings using the procedure in ‘get_embeddings.ipynb’ with modality, organism, and assay tokens assigned according to dataset metadata. The missing assay tokens were replaced with ‘10x 5’ v2’.

#### GeneCompass

GeneCompass [21] is a large-scale cross-species foundation model trained on over 100 million human and mouse cells that encodes gene regulatory programs by combining a transformer encoder for continuous expression vectors with gene-name embeddings enriched by four biological priors (gene families, promoter sequence embeddings, gene-regulatory-network embeddings, and co-expression embeddings). We processed the evaluation datasets using the ‘preprocess.py’ script and obtained the cell embeddings as the CLS-token representations using a custom Python extraction script.

#### scCello

scCello [26] is a transcriptome foundation model based on a transformer architecture augmented with ontology-aware objectives, including masked gene prediction, a cell-type coherence loss, and an ontology alignment loss leveraging the Cell Ontology hierarchy. Pretrained on about 22 million scRNA-seq profiles from the CELLxGENE repository, it learns latent spaces aligned with biological structure. We processed the evaluation datasets using the ‘run_data_transformation.py’ script and extracted the cell embeddings using a custom Python script.

### Evaluation metrics

Integration performance was evaluated using six widely adopted metrics from the scIB framework [8], including three metrics for biological variance conservation and the other three metrics for batch-effect correction. All metrics were computed following the standard scIB implementation.

#### Normalized Mutual Information (NMI)

NMI quantifies the concordance between Leiden-derived clusters and annotated cell-type labels. Higher NMI values indicate better preservation of biological structure, reflecting that cells of the same type cluster together in the embedding space.

#### Adjusted Rand Index (ARI)

ARI measures the clustering agreement between Leiden clusters and cell-type annotations while adjusting for agreement expected by chance. Higher ARI values correspond to more accurate recovery of biologically relevant groupings.

#### Silhouette label

The Silhouette label metric computes the average silhouette width using biological labels. It evaluates how close cells of the same type group together and how effectively they are separated from other types.

Larger values indicate stronger conservation of cell-type structure in the integrated embedding.

#### Batch Removal Adapted Silhouette (BRAS)

BRAS [57] assesses batch mixing within each cell type by computing silhouette values based on batch labels and transforming them such that higher values reflect improved batch mixing. This metric measures the degree to which batch effects are removed without disrupting biological identity.

#### Graph connectivity

Graph connectivity evaluates the structure-preservation by quantifying how well cells of the same biological label remain connected in the k-nearest-neighbor graph. For each cell type, the size of the largest connected component is divided by the total number of cells of that type, and these values are averaged. Scores close to 1 indicate that biological structure is well maintained after integration.

#### scIB score

To summarize overall integration performance, we used the scIB aggregate scoring scheme, which combines metrics related to biological conservation and batch correction. We adopted a 5:5 weighting strategy (biological conservation: batch correction).

### Probing analysis

To assess the types of information encoded in the embeddings produced by each model, we performed a probing analysis, a commonly used approach for interpreting learned embeddings in machine learning [48, 49]. In this framework, a lightweight classifier, referred to as a *probe*, is trained on the embeddings to predict predefined attributes. The underlying assumption is that if a probe achieves high predictive accuracy, the corresponding information is explicitly represented in the embedding space.

In this study, we trained two independent linear probes to predict cell type and batch, respectively. Logistic regression was chosen as the probe classifier due to its low model capacity and reduced tendency to overfit, allowing the evaluation to reflect the information preserved in the embeddings. Probe training and prediction were performed using five-fold cross-validation to ensure robustness.

Probe performance was quantified using accuracy and macro-F1 score.

High cell-type prediction performance indicates strong preservation of biologically meaningful signal, whereas high batch-prediction performance reflects residual technical variation in the embeddings.

### Batch-centralization of embeddings

To examine and mitigate batch-specific biases in the learned embeddings, we applied a batch-centralization procedure as a post-processing step to the embeddings generated by each scFM. Cells were first grouped by batch annotations (e.g., donor, tissue, or disease). For each batch *b*, we estimated its mean embedding vector using at most 10,000 cells. Specifically, if a batch contained more than 10,000 cells, a random subset of 10,000 cells was used to compute 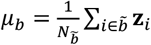, where **z**_*i*_ represents the embedding of cell *i* and 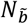 is the number of sampled cells in batch *b*. Each cell embedding was then centered by subtracting the corresponding batch mean, i.e.,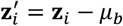.

After centering, we additionally applied L2 normalization to each embedding vector. Unlike more complex integration methods, batch centralization does not modify the relative geometry or variance structure of embeddings, making it a minimal yet interpretable correction that isolates global batch offsets.

## Data availability

All the scRNA-seq data are available from the CELLxGENE database. Specifically, the Liver dataset is available at https://cellxgene.cziscience.com/collections/ff69f0ee-fef6-4895-9f48-6c64a68c8289. The Immune dataset is available at https://cellxgene.cziscience.com/collections/cc431242-35ea-41e1-a100-41e0dec2665b. The HLCA dataset is available at https://cellxgene.cziscience.com/collections/6f6d381a-7701-4781-935c-db10d30de293. The Limb dataset is available at https://cellxgene.cziscience.com/collections/4fefa187-5d14-4f1e-915b-c892ed320aab.

## Acknowledgments

This work has been supported by the National Natural Science Foundation of China (nos. 32341013, 12326614 to S.Z. and 12401661to C.Z.), the Zhejiang Province Vanguard Goose-Leading Initiative (no. 2025C01114), the National Key Research and Development Program of China (nos. 2025YFF1207900 to S.Z. and 2024YFF0729201to C.Z.), and the CAS Project for Young Scientists in Basic Research (no. YSBR-034 to S.Z.).

## Author contributions

Shihua Zhang conceived and supervised the project. Linting Wang and Chihao Zhang implemented the experiments and performed analysis. Linting Wang, Chihao Zhang and Shihua Zhang validated the study. Linting Wang, Chihao Zhang and Shihua Zhang wrote the manuscript. All authors read and approved the final manuscript.

