## Supplemental Tables and Figures for "Batch Effects Remain a Fundamental Barrier to Universal Embeddings in Single-Cell Foundation Models"

**a**

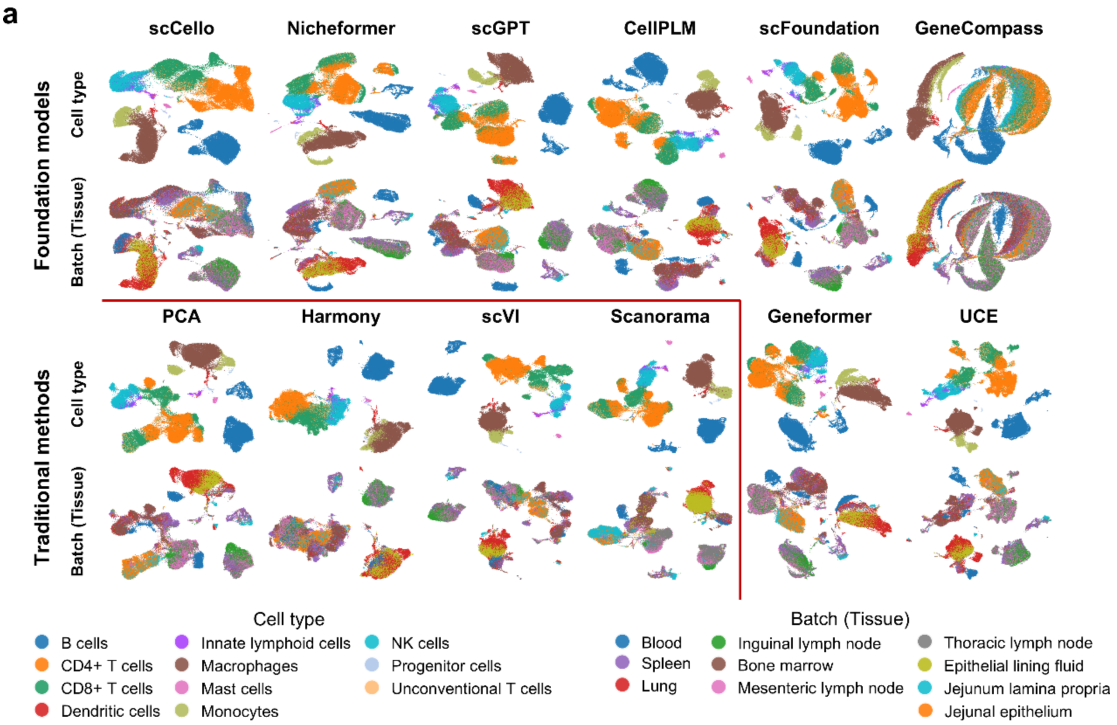


**b**

| | Bio conservation | | | Batch correction | | Aggregate score | | |
|---|---|---|---|---|---|---|---|---|
| Method | Leiden NMI | Leiden ARI | Silhouette label | BRAS | Graph connectivity | Bio conservation | Batch correction | Total |
| scVI | 0.79 | 0.74 | 0.53 | 0.82 | 0.87 | 0.69 | 0.85 | 0.77 |
| scCello | 0.79 | 0.70 | 0.64 | 0.71 | 0.90 | 0.71 | 0.81 | 0.76 |
| Harmony | 0.74 | 0.69 | 0.55 | 0.86 | 0.78 | 0.66 | 0.82 | 0.74 |
| Scanorama | 0.76 | 0.68 | 0.56 | 0.77 | 0.79 | 0.67 | 0.78 | 0.73 |
| PCA | 0.74 | 0.65 | 0.59 | 0.67 | 0.85 | 0.66 | 0.76 | 0.71 |
| scGPT | 0.71 | 0.62 | 0.57 | 0.66 | 0.87 | 0.63 | 0.77 | 0.70 |
| CellPLM | 0.70 | 0.60 | 0.57 | 0.67 | 0.86 | 0.62 | 0.77 | 0.69 |
| Geneformer | 0.68 | 0.58 | 0.53 | 0.77 | 0.79 | 0.59 | 0.78 | 0.69 |
| scFoundation | 0.73 | 0.64 | 0.58 | 0.63 | 0.80 | 0.65 | 0.71 | 0.68 |
| Nicheformer | 0.68 | 0.49 | 0.58 | 0.71 | 0.82 | 0.58 | 0.77 | 0.67 |
| UCE | 0.71 | 0.57 | 0.58 | 0.68 | 0.56 | 0.62 | 0.62 | 0.62 |
| GeneCompass | 0.53 | 0.36 | 0.46 | 0.78 | 0.80 | 0.45 | 0.79 | 0.62 |

**Figure S1. Evaluation of batch effects on the Immune dataset with tissue as the batch source. a**, UMAP visualizations of cell embeddings derived from the traditional integration methods and scFMs. **b**, Detailed scIB scores of traditional methods and scFMs.

**a**

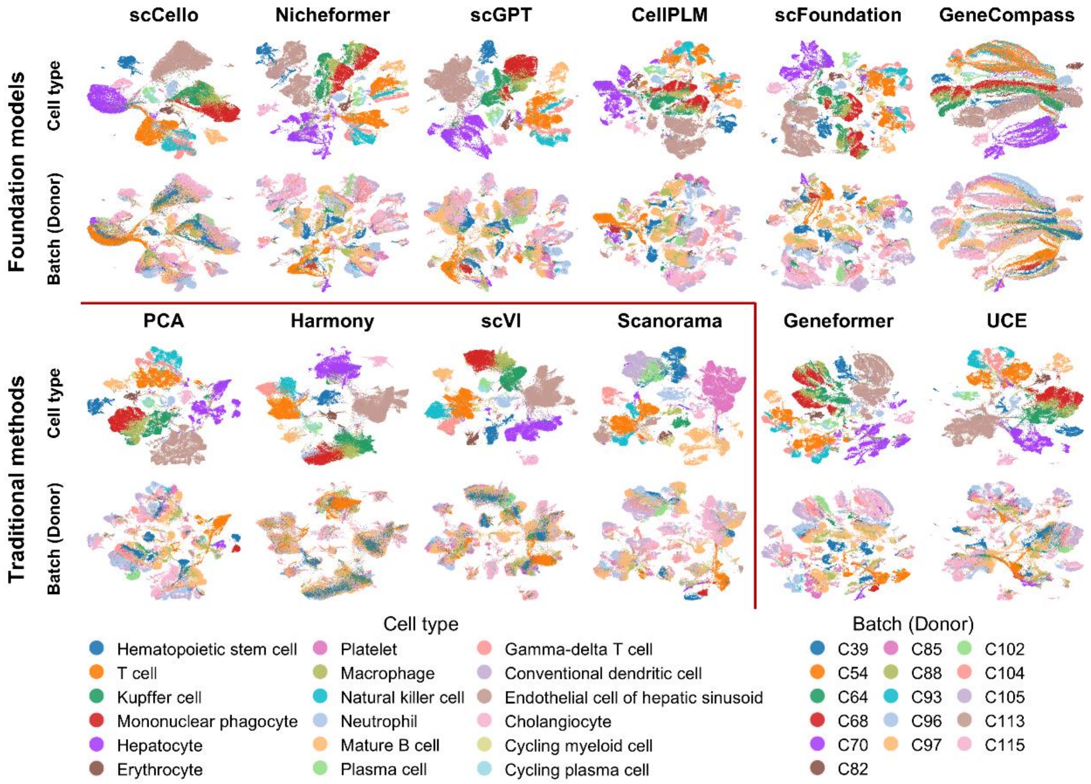


**b**

| Method | Bio conservation: Leiden NMI | Bio conservation: Leiden ARI | Bio conservation: Silhouette label | Batch correction: BRAS | Batch correction: Graph connectivity | Aggregate score: Bio conservation | Aggregate score: Batch correction | Aggregate score: Total |
|---|---|---|---|---|---|---|---|---|
| **scVI** | 0.82 | 0.72 | 0.57 | 0.79 | 0.92 | 0.70 | 0.86 | 0.78 |
| **Harmony** | 0.83 | 0.82 | 0.60 | 0.81 | 0.81 | 0.75 | 0.81 | 0.78 |
| **scCello** | 0.78 | 0.72 | 0.62 | 0.63 | 0.87 | 0.71 | 0.75 | 0.73 |
| **Scanorama** | 0.74 | 0.55 | 0.59 | 0.70 | 0.89 | 0.63 | 0.79 | 0.71 |
| **scGPT** | 0.74 | 0.66 | 0.58 | 0.62 | 0.90 | 0.66 | 0.76 | 0.71 |
| **UCE** | 0.77 | 0.73 | 0.59 | 0.70 | 0.67 | 0.70 | 0.68 | 0.69 |
| **PCA** | 0.76 | 0.54 | 0.60 | 0.59 | 0.86 | 0.63 | 0.73 | 0.68 |
| **Nicheformer** | 0.68 | 0.48 | 0.58 | 0.62 | 0.87 | 0.58 | 0.75 | 0.66 |
| **Geneformer** | 0.63 | 0.40 | 0.52 | 0.67 | 0.87 | 0.52 | 0.77 | 0.64 |
| **scFoundation** | 0.65 | 0.39 | 0.58 | 0.58 | 0.89 | 0.54 | 0.74 | 0.64 |
| **CellPLM** | 0.64 | 0.38 | 0.58 | 0.53 | 0.90 | 0.54 | 0.72 | 0.63 |
| **GeneCompass** | 0.58 | 0.34 | 0.41 | 0.63 | 0.86 | 0.44 | 0.74 | 0.59 |

**Figure S2. Evaluation of batch effects on the Liver dataset with the donor as the batch source. a**, UMAP visualizations of cell embeddings derived from the traditional integration methods and scFMs. **b**, Detailed scIB scores of traditional methods and scFMs.

**a**

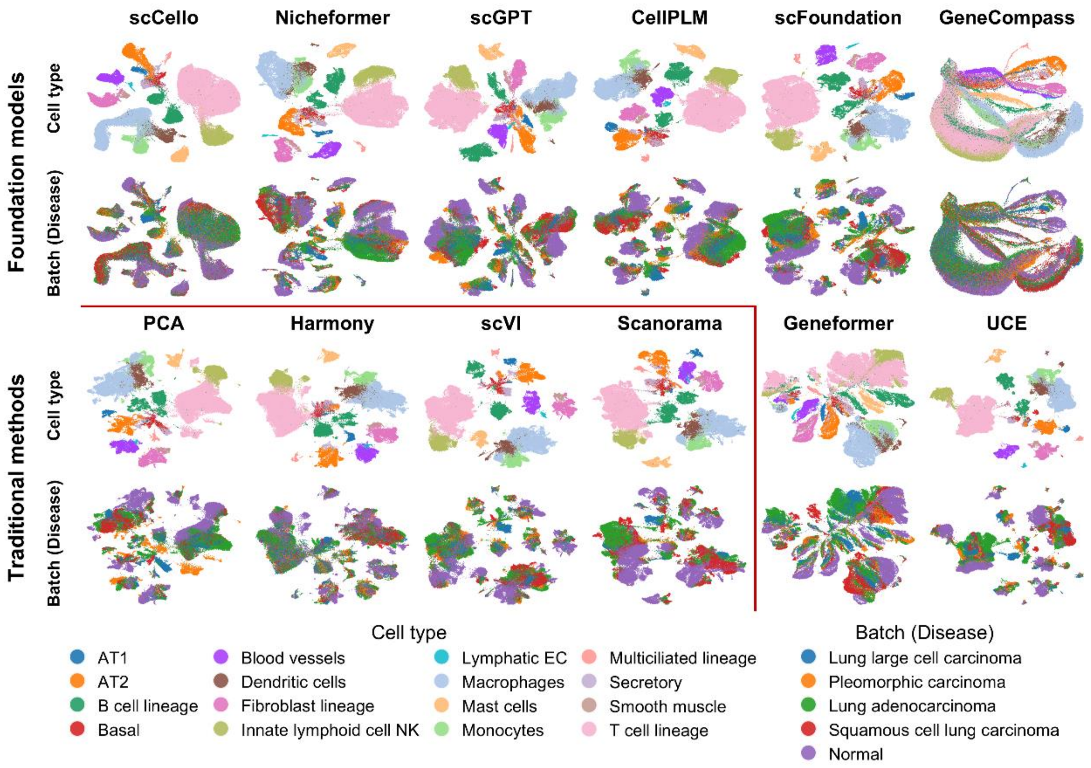


**b**

| Method | Bio conservation: Leiden NMI | Bio conservation: Leiden ARI | Bio conservation: Silhouette label | Batch correction: BRAS | Batch correction: Graph connectivity | Aggregate score: Bio conservation | Aggregate score: Batch correction | Aggregate score: Total |
|---|---|---|---|---|---|---|---|---|
| **Harmony** | 0.85 | 0.86 | 0.58 | 0.86 | 0.88 | 0.76 | 0.87 | 0.82 |
| **scVI** | 0.85 | 0.81 | 0.55 | 0.82 | 0.95 | 0.74 | 0.89 | 0.81 |
| **scGPT** | 0.85 | 0.86 | 0.56 | 0.77 | 0.96 | 0.76 | 0.86 | 0.81 |
| **Nicheformer** | 0.85 | 0.82 | 0.56 | 0.78 | 0.93 | 0.75 | 0.85 | 0.80 |
| **CellPLM** | 0.83 | 0.81 | 0.56 | 0.74 | 0.95 | 0.74 | 0.85 | 0.79 |
| **Scanorama** | 0.81 | 0.71 | 0.56 | 0.78 | 0.93 | 0.69 | 0.85 | 0.77 |
| **scCello** | 0.81 | 0.60 | 0.72 | 0.74 | 0.92 | 0.71 | 0.83 | 0.77 |
| **scFoundation** | 0.80 | 0.64 | 0.57 | 0.74 | 0.95 | 0.67 | 0.85 | 0.76 |
| **PCA** | 0.81 | 0.74 | 0.59 | 0.71 | 0.88 | 0.71 | 0.79 | 0.75 |
| **Geneformer** | 0.66 | 0.43 | 0.51 | 0.78 | 0.90 | 0.53 | 0.84 | 0.69 |
| **UCE** | 0.83 | 0.80 | 0.56 | 0.78 | 0.40 | 0.73 | 0.59 | 0.66 |
| **GeneCompass** | 0.56 | 0.26 | 0.46 | 0.74 | 0.89 | 0.43 | 0.81 | 0.62 |

**Figure S3. Evaluation of batch effects on the HLCA dataset with disease as the batch source. a**, UMAP visualizations of cell embeddings derived from the traditional integration methods and scFMs. **b**, Detailed scIB scores of traditional methods and scFMs.

**a**

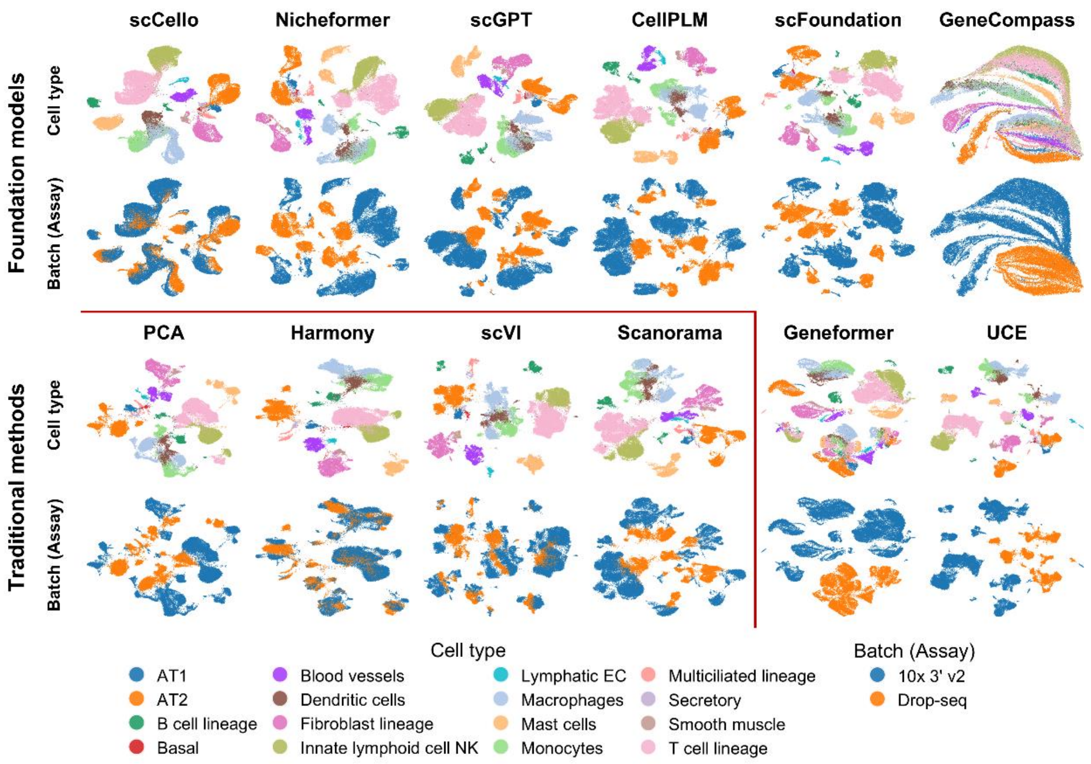


**b**

| Method | Bio conservation: Leiden NMI | Bio conservation: Leiden ARI | Bio conservation: Silhouette label | Batch correction: BRAS | Batch correction: Graph connectivity | Aggregate score: Bio conservation | Aggregate score: Batch correction | Aggregate score: Total |
|---|---|---|---|---|---|---|---|---|
| scVI | 0.87 | 0.81 | 0.55 | 0.86 | 0.94 | 0.75 | 0.90 | 0.82 |
| Harmony | 0.85 | 0.79 | 0.59 | 0.87 | 0.87 | 0.74 | 0.87 | 0.81 |
| scCello | 0.85 | 0.78 | 0.70 | 0.65 | 0.92 | 0.78 | 0.78 | 0.78 |
| Scanorama | 0.80 | 0.71 | 0.55 | 0.59 | 0.90 | 0.68 | 0.74 | 0.71 |
| scGPT | 0.79 | 0.66 | 0.56 | 0.57 | 0.93 | 0.67 | 0.75 | 0.71 |
| PCA | 0.77 | 0.61 | 0.58 | 0.60 | 0.90 | 0.65 | 0.75 | 0.70 |
| CellPLM | 0.80 | 0.66 | 0.57 | 0.49 | 0.95 | 0.68 | 0.72 | 0.70 |
| scFoundation | 0.76 | 0.58 | 0.57 | 0.54 | 0.94 | 0.64 | 0.74 | 0.69 |
| Nicheformer | 0.77 | 0.56 | 0.56 | 0.49 | 0.92 | 0.63 | 0.71 | 0.67 |
| UCE | 0.79 | 0.64 | 0.55 | 0.68 | 0.32 | 0.66 | 0.50 | 0.58 |
| Geneformer | 0.63 | 0.29 | 0.47 | 0.43 | 0.69 | 0.46 | 0.56 | 0.51 |
| GeneCompass | 0.54 | 0.20 | 0.45 | 0.44 | 0.81 | 0.40 | 0.63 | 0.51 |

**Figure S4. Evaluation of batch effects on the HLCA dataset with assay as the batch source. a**, UMAP visualizations of cell embeddings derived from the traditional integration methods and scFMs. **b**, Detailed scIB scores of traditional methods and scFMs.

a

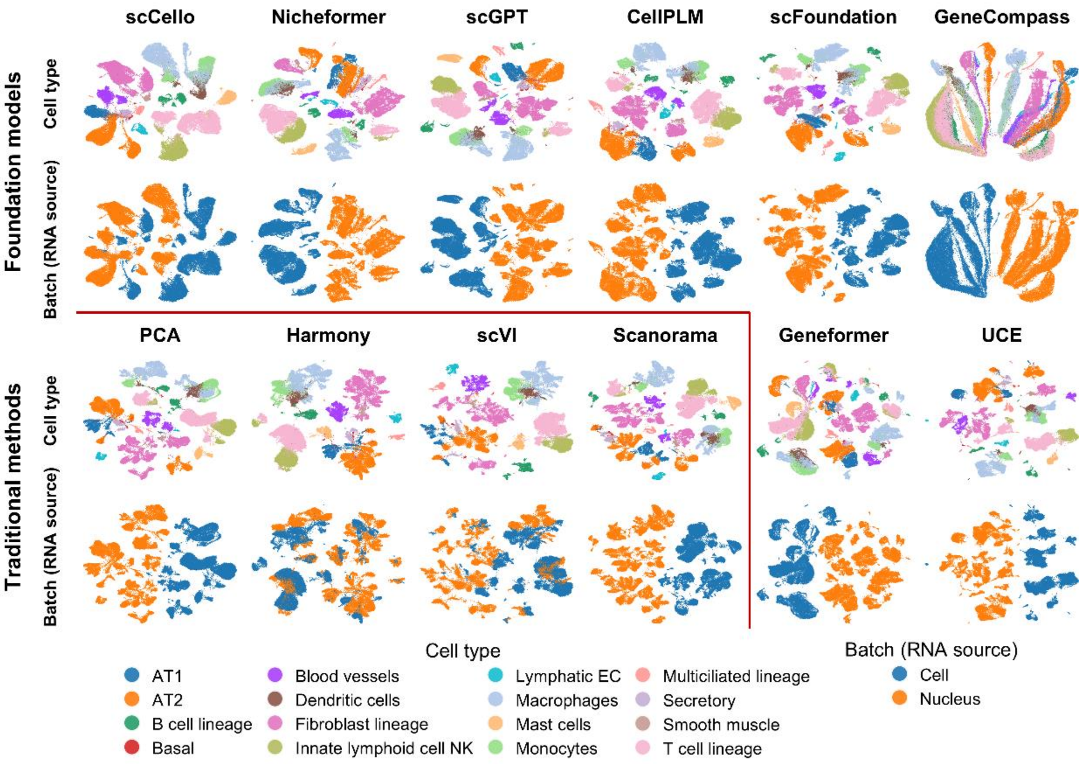


b

| Method | Bio conservation: Leiden NMI | Bio conservation: Leiden ARI | Bio conservation: Silhouette label | Batch correction: BRAS | Batch correction: Graph connectivity | Aggregate score: Bio conservation | Aggregate score: Batch correction | Aggregate score: Total |
|---|---|---|---|---|---|---|---|---|
| **scVI** | 0.84 | 0.75 | 0.55 | 0.83 | 0.89 | 0.71 | 0.86 | 0.79 |
| **Harmony** | 0.82 | 0.67 | 0.61 | 0.84 | 0.86 | 0.70 | 0.85 | 0.78 |
| **scCello** | 0.80 | 0.64 | 0.69 | 0.56 | 0.88 | 0.71 | 0.72 | 0.72 |
| **scFoundation** | 0.80 | 0.65 | 0.57 | 0.39 | 0.78 | 0.67 | 0.58 | 0.63 |
| **PCA** | 0.76 | 0.54 | 0.59 | 0.38 | 0.82 | 0.63 | 0.60 | 0.62 |
| **CellPLM** | 0.78 | 0.63 | 0.56 | 0.33 | 0.79 | 0.66 | 0.56 | 0.61 |
| **scGPT** | 0.79 | 0.61 | 0.54 | 0.30 | 0.80 | 0.64 | 0.55 | 0.60 |
| **GeneCompass** | 0.59 | 0.40 | 0.45 | 0.60 | 0.76 | 0.48 | 0.68 | 0.58 |
| **Nicheformer** | 0.79 | 0.63 | 0.54 | 0.28 | 0.72 | 0.65 | 0.50 | 0.58 |
| **UCE** | 0.76 | 0.54 | 0.57 | 0.50 | 0.41 | 0.62 | 0.46 | 0.54 |
| **Geneformer** | 0.68 | 0.43 | 0.49 | 0.43 | 0.65 | 0.53 | 0.54 | 0.54 |
| **Scanorama** | 0.74 | 0.50 | 0.50 | 0.18 | 0.75 | 0.58 | 0.46 | 0.52 |

**Figure S5. Evaluation of batch effects on the HLCA dataset with RNA source as the batch source. a**, UMAP visualizations of cell embeddings derived from the traditional integration methods and scFMs. **b**, Detailed scIB scores of traditional methods and scFMs.

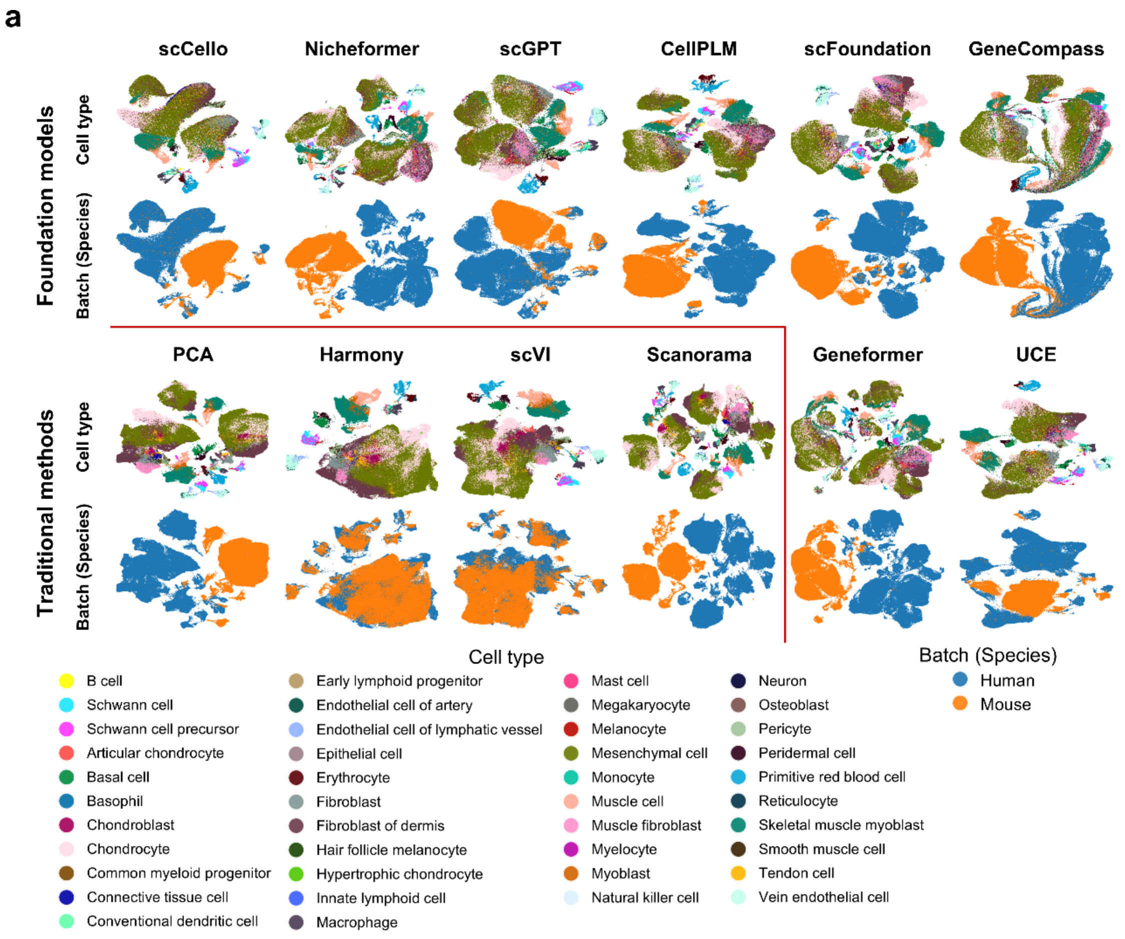


**b**

| Method | Bio conservation: Leiden NMI | Bio conservation: Leiden ARI | Bio conservation: Silhouette label | Batch correction: BRAS | Batch correction: Graph connectivity | Aggregate score: Bio conservation | Aggregate score: Batch correction | Aggregate score: Total |
|---|---|---|---|---|---|---|---|---|
| scVI | 0.50 | 0.19 | 0.51 | 0.77 | 0.85 | 0.40 | 0.81 | 0.60 |
| Harmony | 0.51 | 0.19 | 0.51 | 0.81 | 0.72 | 0.40 | 0.77 | 0.58 |
| scGPT | 0.45 | 0.22 | 0.48 | 0.62 | 0.76 | 0.38 | 0.69 | 0.54 |
| scFoundation | 0.44 | 0.21 | 0.50 | 0.62 | 0.75 | 0.38 | 0.68 | 0.53 |
| UCE | 0.46 | 0.23 | 0.50 | 0.69 | 0.62 | 0.40 | 0.65 | 0.53 |
| scCello | 0.42 | 0.18 | 0.46 | 0.62 | 0.67 | 0.35 | 0.65 | 0.50 |
| PCA | 0.47 | 0.11 | 0.50 | 0.61 | 0.65 | 0.36 | 0.63 | 0.49 |
| CellPLM | 0.40 | 0.11 | 0.48 | 0.59 | 0.68 | 0.33 | 0.64 | 0.49 |
| Geneformer | 0.35 | 0.06 | 0.47 | 0.63 | 0.59 | 0.29 | 0.61 | 0.45 |
| GeneCompass | 0.31 | 0.08 | 0.44 | 0.58 | 0.53 | 0.28 | 0.56 | 0.42 |
| Scanorama | 0.44 | 0.09 | 0.45 | 0.40 | 0.60 | 0.32 | 0.50 | 0.41 |
| Nicheformer | 0.38 | 0.11 | 0.45 | 0.43 | 0.59 | 0.31 | 0.51 | 0.41 |

**Figure S6. Evaluation of batch effects on the limb dataset with species as the batch source. a**, UMAP visualizations of cell embeddings derived from the traditional integration methods and scFMs. **b**, Detailed scIB scores of traditional methods and scFMs.

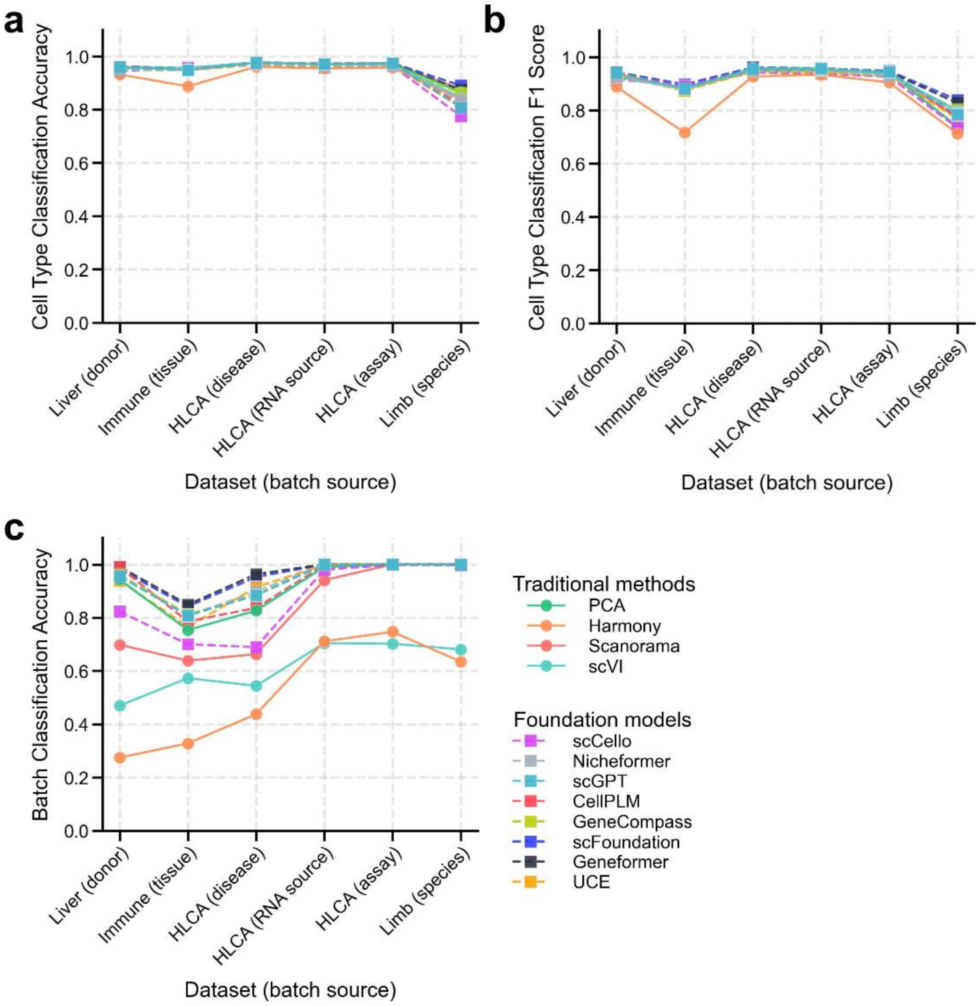


**Figure S7. Evaluation of cell-type or batch information retained within the embeddings generated from traditional methods and scFMs. a** and **b**, Cell-type classification accuracy (**a**) and macro-F1 score (**b**) across six datasets of traditional methods and scFMs. **c**, Batch classification accuracy across six datasets of traditional methods and scFMs.

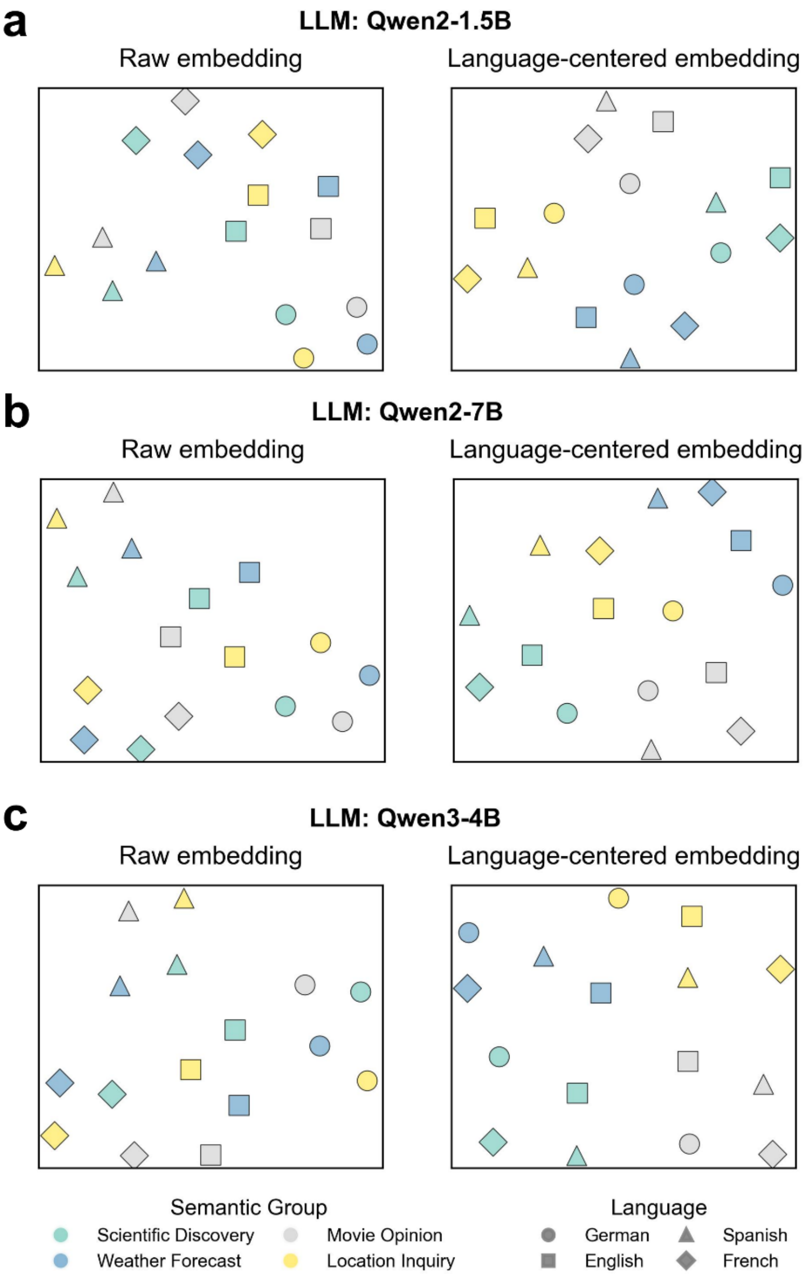


**Figure S8. The visualizations of the sentence embedding space in the open-source LLM Qwen. a-c**, UMAP visualizations of raw and language-centered sentence embeddings generated by Qwen2-1.5B (**a**), Qwen2-7B (**b**), and Qwen3-4B (**c**) on a multilingual corpus, illustrating semantic alignment through language centralization.

**a**

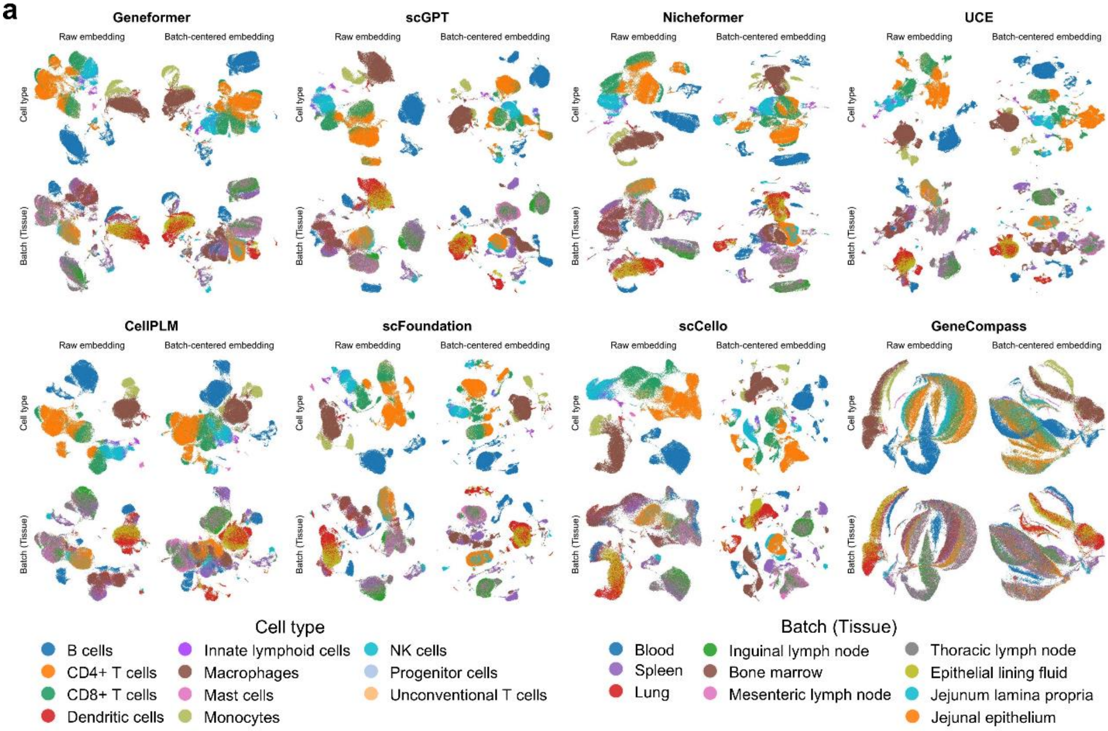


**b**

| | Bio conservation | | | Batch correction | | Aggregate score | | |
|---|---|---|---|---|---|---|---|---|
| **Method** | Leiden NMI | Leiden ARI | Silhouette label | BRAS | Graph connectivity | Bio conservation | Batch correction | Total |
| **scVI** | 0.79 | 0.74 | 0.53 | 0.82 | 0.87 | 0.69 | 0.85 | 0.77 |
| **scCello** | 0.79 | 0.70 | 0.64 | 0.71 | 0.90 | 0.71 | 0.81 | 0.76 |
| **Harmony** | 0.74 | 0.69 | 0.55 | 0.86 | 0.78 | 0.66 | 0.82 | 0.74 |
| **Scanorama** | 0.76 | 0.68 | 0.56 | 0.77 | 0.79 | 0.67 | 0.78 | 0.73 |
| **PCA** | 0.74 | 0.65 | 0.59 | 0.67 | 0.85 | 0.66 | 0.76 | 0.71 |
| **scGPT** | 0.71 | 0.62 | 0.57 | 0.66 | 0.87 | 0.63 | 0.77 | 0.70 |
| **CellPLM** | 0.70 | 0.60 | 0.57 | 0.67 | 0.86 | 0.62 | 0.77 | 0.69 |
| **Geneformer** | 0.68 | 0.58 | 0.53 | 0.77 | 0.79 | 0.59 | 0.78 | 0.69 |
| **scFoundation** | 0.73 | 0.64 | 0.58 | 0.63 | 0.80 | 0.65 | 0.71 | 0.68 |
| **Nicheformer** | 0.68 | 0.49 | 0.58 | 0.71 | 0.82 | 0.58 | 0.77 | 0.67 |
| **Geneformer_batch** | 0.63 | 0.55 | 0.50 | 0.77 | 0.74 | 0.56 | 0.76 | 0.66 |
| **scGPT_batch** | 0.64 | 0.52 | 0.54 | 0.67 | 0.74 | 0.57 | 0.71 | 0.64 |
| **scFoundation_batch** | 0.65 | 0.55 | 0.53 | 0.65 | 0.72 | 0.58 | 0.68 | 0.63 |
| **CellPLM_batch** | 0.61 | 0.51 | 0.51 | 0.65 | 0.77 | 0.54 | 0.71 | 0.63 |
| **UCE** | 0.71 | 0.57 | 0.58 | 0.68 | 0.56 | 0.62 | 0.62 | 0.62 |
| **GeneCompass** | 0.53 | 0.36 | 0.46 | 0.78 | 0.80 | 0.45 | 0.79 | 0.62 |
| **scCello_batch** | 0.67 | 0.57 | 0.58 | 0.51 | 0.69 | 0.60 | 0.60 | 0.60 |
| **UCE_batch** | 0.66 | 0.55 | 0.53 | 0.69 | 0.56 | 0.58 | 0.63 | 0.60 |
| **Nicheformer_batch** | 0.62 | 0.48 | 0.54 | 0.66 | 0.63 | 0.55 | 0.65 | 0.60 |
| **GeneCompass_batch** | 0.52 | 0.38 | 0.44 | 0.80 | 0.67 | 0.45 | 0.74 | 0.59 |

**Figure S9. Comparison of embedding from scFMs before and after batch-centralization on the immune dataset. a**, Comparison of UMAP visualizations of raw and batch-centered embeddings derived from scFMs. **b**, Detailed scIB scores of traditional methods and scFMs.

**a**

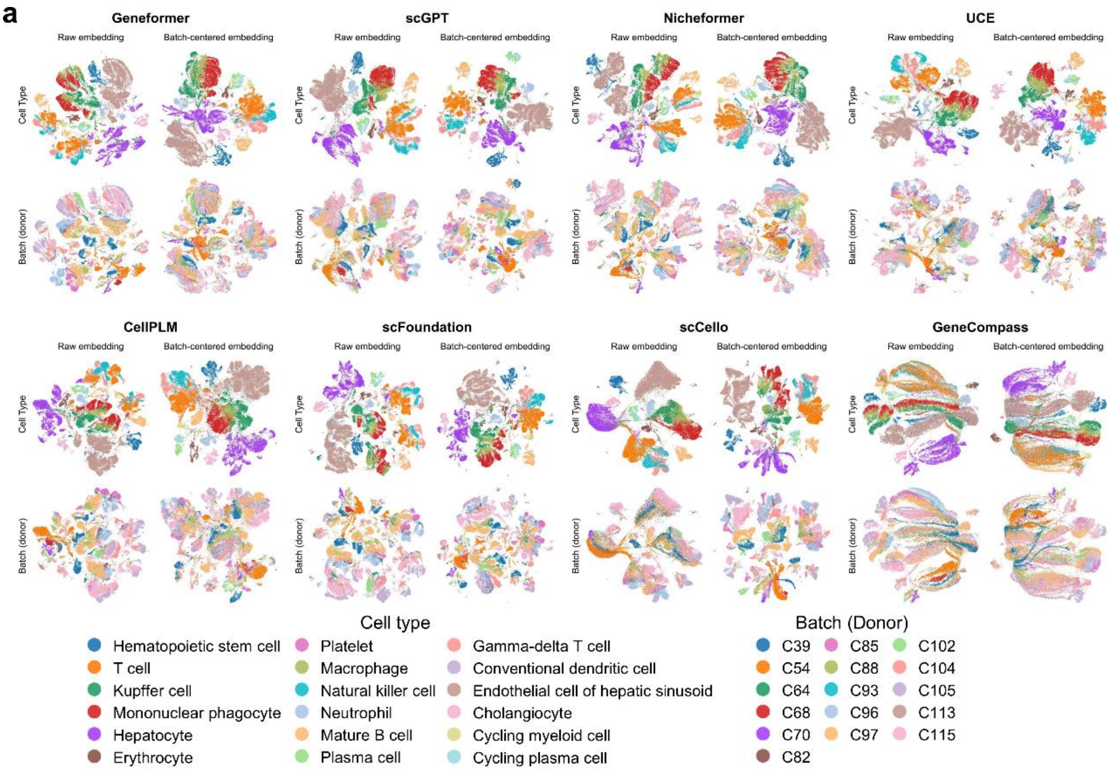


**b**

| Method | Bio conservation | | | Batch correction | | Aggregate score | | |
|---|---|---|---|---|---|---|---|---|
| | Leiden NMI | Leiden ARI | Silhouette label | BRAS | Graph connectivity | Bio conservation | Batch correction | Total |
| scVI | 0.82 | 0.72 | 0.57 | 0.79 | 0.92 | 0.70 | 0.86 | 0.78 |
| Harmony | 0.83 | 0.82 | 0.60 | 0.81 | 0.81 | 0.75 | 0.81 | 0.78 |
| CellPLM_batch | 0.74 | 0.66 | 0.56 | 0.70 | 0.92 | 0.65 | 0.81 | 0.73 |
| scCello | 0.78 | 0.72 | 0.62 | 0.63 | 0.87 | 0.71 | 0.75 | 0.73 |
| Geneformer_batch | 0.73 | 0.64 | 0.51 | 0.79 | 0.87 | 0.63 | 0.83 | 0.73 |
| scGPT_batch | 0.73 | 0.60 | 0.56 | 0.73 | 0.88 | 0.63 | 0.81 | 0.72 |
| UCE_batch | 0.80 | 0.74 | 0.57 | 0.80 | 0.65 | 0.70 | 0.72 | 0.71 |
| Scanorama | 0.74 | 0.55 | 0.59 | 0.70 | 0.89 | 0.63 | 0.79 | 0.71 |
| scGPT | 0.74 | 0.66 | 0.58 | 0.62 | 0.90 | 0.66 | 0.76 | 0.71 |
| Nicheformer_batch | 0.72 | 0.57 | 0.56 | 0.73 | 0.82 | 0.62 | 0.78 | 0.70 |
| UCE | 0.77 | 0.73 | 0.59 | 0.70 | 0.67 | 0.70 | 0.68 | 0.69 |
| scFoundation_batch | 0.67 | 0.44 | 0.56 | 0.73 | 0.90 | 0.56 | 0.82 | 0.69 |
| PCA | 0.76 | 0.54 | 0.60 | 0.59 | 0.86 | 0.63 | 0.73 | 0.68 |
| Nicheformer | 0.68 | 0.48 | 0.58 | 0.62 | 0.87 | 0.58 | 0.75 | 0.66 |
| scCello_batch | 0.69 | 0.47 | 0.60 | 0.68 | 0.78 | 0.58 | 0.73 | 0.66 |
| Geneformer | 0.63 | 0.40 | 0.52 | 0.67 | 0.87 | 0.52 | 0.77 | 0.64 |
| scFoundation | 0.65 | 0.39 | 0.58 | 0.58 | 0.89 | 0.54 | 0.74 | 0.64 |
| CellPLM | 0.64 | 0.38 | 0.58 | 0.53 | 0.90 | 0.54 | 0.72 | 0.63 |
| GeneCompass_batch | 0.60 | 0.38 | 0.41 | 0.75 | 0.82 | 0.46 | 0.78 | 0.62 |
| GeneCompass | 0.58 | 0.34 | 0.41 | 0.63 | 0.86 | 0.44 | 0.74 | 0.59 |

**Figure S10. Comparison of embedding from scFMs before and after batch-centralization on the liver dataset. a**, Comparison of UMAP visualizations of raw and batch-centered embeddings derived from scFMs. **b**, Detailed scIB scores of traditional methods and scFMs.

**a**

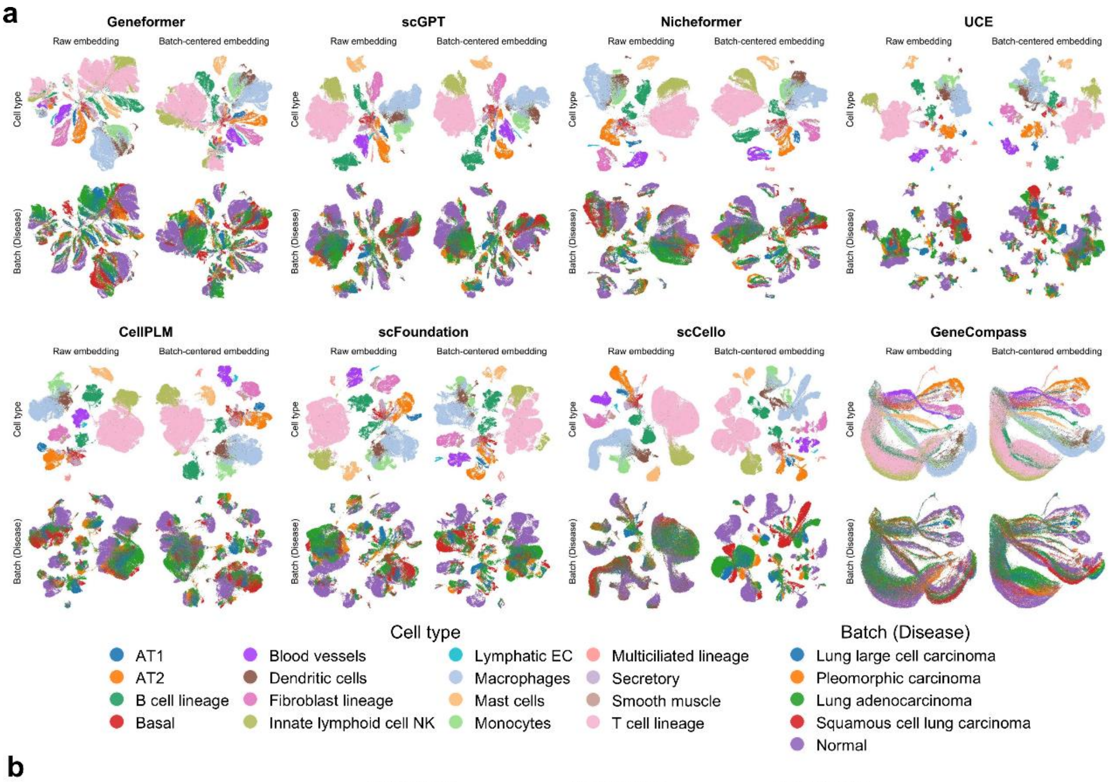


**b**

| | Bio conservation | | | Batch correction | | Aggregate score | | |
|---|---|---|---|---|---|---|---|---|
| Method | Leiden NMI | Leiden ARI | Silhouette label | BRAS | Graph connectivity | Bio conservation | Batch correction | Total |
| Harmony | 0.85 | 0.86 | 0.58 | 0.86 | 0.88 | 0.76 | 0.87 | 0.82 |
| scVI | 0.85 | 0.81 | 0.55 | 0.82 | 0.95 | 0.74 | 0.89 | 0.81 |
| scGPT | 0.85 | 0.86 | 0.56 | 0.77 | 0.96 | 0.76 | 0.86 | 0.81 |
| CellPLM_batch | 0.85 | 0.86 | 0.54 | 0.78 | 0.95 | 0.75 | 0.86 | 0.81 |
| Nicheformer | 0.85 | 0.82 | 0.56 | 0.78 | 0.93 | 0.75 | 0.85 | 0.80 |
| scGPT_batch | 0.83 | 0.75 | 0.55 | 0.81 | 0.96 | 0.71 | 0.88 | 0.80 |
| CellPLM | 0.83 | 0.81 | 0.56 | 0.74 | 0.95 | 0.74 | 0.85 | 0.79 |
| Nicheformer_batch | 0.81 | 0.71 | 0.56 | 0.80 | 0.92 | 0.69 | 0.86 | 0.78 |
| Scanorama | 0.81 | 0.71 | 0.56 | 0.78 | 0.93 | 0.69 | 0.85 | 0.77 |
| scCello | 0.81 | 0.60 | 0.72 | 0.74 | 0.92 | 0.71 | 0.83 | 0.77 |
| scFoundation | 0.80 | 0.64 | 0.57 | 0.74 | 0.95 | 0.67 | 0.85 | 0.76 |
| scFoundation_batch | 0.77 | 0.63 | 0.56 | 0.78 | 0.95 | 0.65 | 0.86 | 0.76 |
| PCA | 0.81 | 0.74 | 0.59 | 0.71 | 0.88 | 0.71 | 0.79 | 0.75 |
| UCE_batch | 0.80 | 0.83 | 0.55 | 0.82 | 0.56 | 0.73 | 0.69 | 0.71 |
| Geneformer_batch | 0.64 | 0.49 | 0.50 | 0.82 | 0.92 | 0.55 | 0.87 | 0.71 |
| scCello_batch | 0.73 | 0.41 | 0.68 | 0.70 | 0.91 | 0.61 | 0.80 | 0.70 |
| Geneformer | 0.66 | 0.43 | 0.51 | 0.78 | 0.90 | 0.53 | 0.84 | 0.69 |
| UCE | 0.83 | 0.80 | 0.56 | 0.78 | 0.40 | 0.73 | 0.59 | 0.66 |
| GeneCompass_batch | 0.54 | 0.38 | 0.46 | 0.78 | 0.85 | 0.46 | 0.82 | 0.64 |
| GeneCompass | 0.56 | 0.26 | 0.46 | 0.74 | 0.89 | 0.43 | 0.81 | 0.62 |

**Figure S11. Comparison of embedding from scFMs before and after batch-centralization on the HLCA dataset with disease as the batch source. a**, Comparison of UMAP visualizations of raw and batch-centered embeddings derived from scFMs. **b**, Detailed scIB scores of traditional methods and scFMs.

**a**

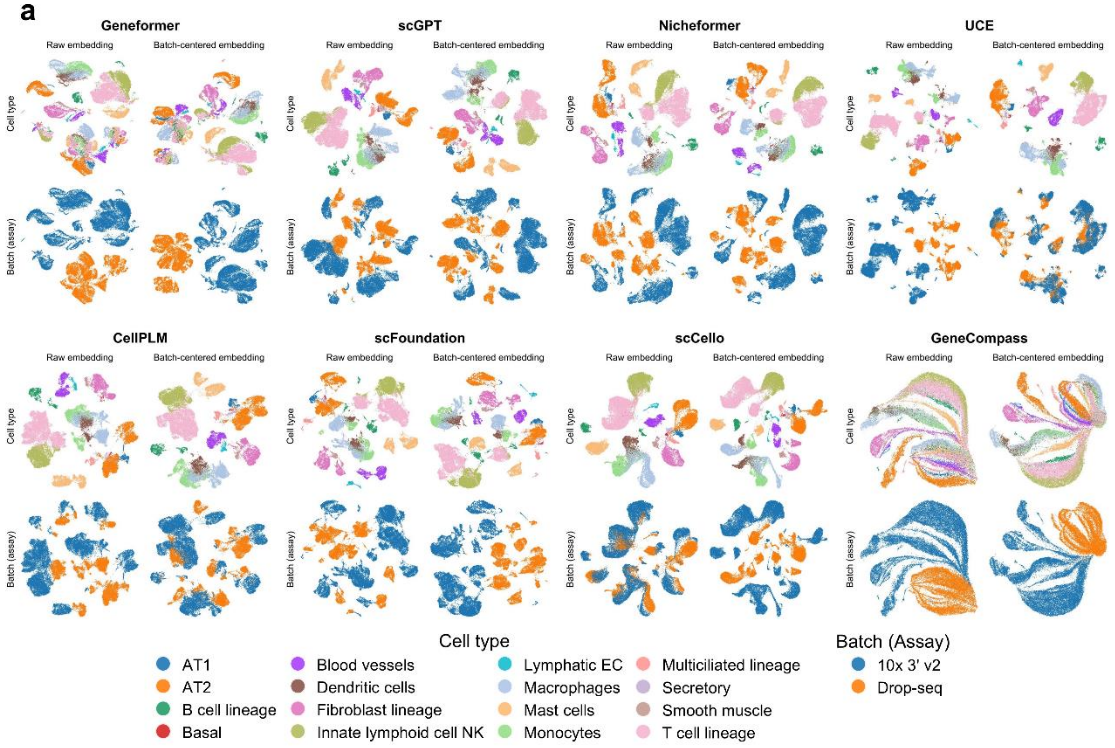


**b**

| Method | Bio conservation | | | Batch correction | | Aggregate score | | |
|---|---|---|---|---|---|---|---|---|
| | Leiden NMI | Leiden ARI | Silhouette label | BRAS | Graph connectivity | Bio conservation | Batch correction | Total |
| scVI | 0.87 | 0.81 | 0.55 | 0.86 | 0.94 | 0.75 | 0.90 | 0.82 |
| Harmony | 0.85 | 0.79 | 0.59 | 0.87 | 0.87 | 0.74 | 0.87 | 0.81 |
| scCello | 0.85 | 0.78 | 0.70 | 0.65 | 0.92 | 0.78 | 0.78 | 0.78 |
| CellPLM_batch | 0.82 | 0.76 | 0.57 | 0.73 | 0.95 | 0.72 | 0.84 | 0.78 |
| scGPT_batch | 0.80 | 0.65 | 0.57 | 0.77 | 0.93 | 0.67 | 0.85 | 0.76 |
| Nicheformer_batch | 0.78 | 0.64 | 0.58 | 0.76 | 0.80 | 0.67 | 0.78 | 0.72 |
| scFoundation_batch | 0.76 | 0.59 | 0.58 | 0.75 | 0.84 | 0.64 | 0.80 | 0.72 |
| scCello_batch | 0.80 | 0.65 | 0.69 | 0.64 | 0.80 | 0.71 | 0.72 | 0.72 |
| Scanorama | 0.80 | 0.71 | 0.55 | 0.59 | 0.90 | 0.68 | 0.74 | 0.71 |
| scGPT | 0.79 | 0.66 | 0.56 | 0.57 | 0.93 | 0.67 | 0.75 | 0.71 |
| PCA | 0.77 | 0.61 | 0.58 | 0.60 | 0.90 | 0.65 | 0.75 | 0.70 |
| CellPLM | 0.80 | 0.66 | 0.57 | 0.49 | 0.95 | 0.68 | 0.72 | 0.70 |
| scFoundation | 0.76 | 0.58 | 0.57 | 0.54 | 0.94 | 0.64 | 0.74 | 0.69 |
| UCE_batch | 0.87 | 0.83 | 0.56 | 0.79 | 0.44 | 0.75 | 0.62 | 0.68 |
| Nicheformer | 0.77 | 0.56 | 0.56 | 0.49 | 0.92 | 0.63 | 0.71 | 0.67 |
| Geneformer_batch | 0.62 | 0.28 | 0.52 | 0.80 | 0.69 | 0.47 | 0.75 | 0.61 |
| UCE | 0.79 | 0.64 | 0.55 | 0.68 | 0.32 | 0.66 | 0.50 | 0.58 |
| GeneCompass_batch | 0.54 | 0.25 | 0.48 | 0.74 | 0.61 | 0.42 | 0.68 | 0.55 |
| Geneformer | 0.63 | 0.29 | 0.47 | 0.43 | 0.69 | 0.46 | 0.56 | 0.51 |
| GeneCompass | 0.54 | 0.20 | 0.45 | 0.44 | 0.81 | 0.40 | 0.63 | 0.51 |

**Figure S12. Comparison of embedding from scFMs before and after batch-centralization on the HLCA dataset with assay as batch source. a**, Comparison of UMAP visualizations of raw and batch-centered embeddings derived from scFMs. **b**, Detailed scIB scores of traditional methods and scFMs.

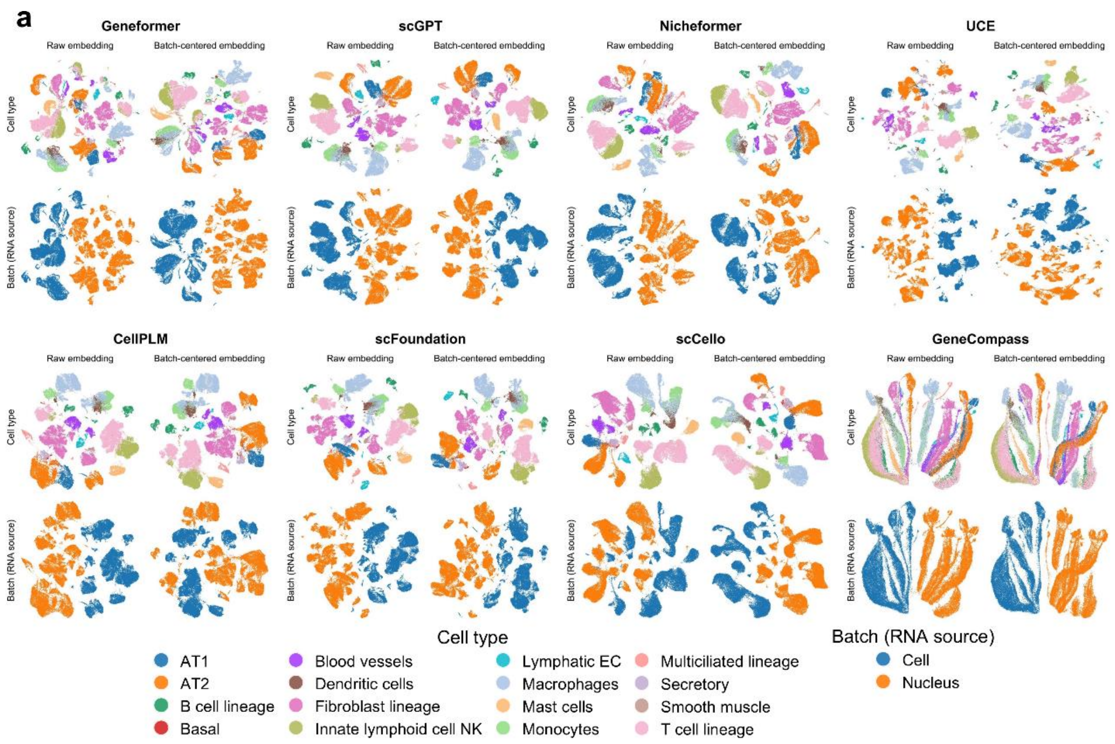


**b**

| Method | Bio conservation: Leiden NMI | Bio conservation: Leiden ARI | Bio conservation: Silhouette label | Batch correction: BRAS | Batch correction: Graph connectivity | Aggregate score: Bio conservation | Aggregate score: Batch correction | Aggregate score: Total |
|---|---|---|---|---|---|---|---|---|
| scVI | 0.84 | 0.75 | 0.55 | 0.83 | 0.89 | 0.71 | 0.86 | 0.79 |
| Harmony | 0.82 | 0.67 | 0.61 | 0.84 | 0.86 | 0.70 | 0.85 | 0.78 |
| CellPLM_batch | 0.79 | 0.64 | 0.58 | 0.66 | 0.90 | 0.67 | 0.78 | 0.72 |
| scCello | 0.80 | 0.64 | 0.69 | 0.56 | 0.88 | 0.71 | 0.72 | 0.72 |
| scFoundation_batch | 0.78 | 0.59 | 0.59 | 0.71 | 0.81 | 0.65 | 0.76 | 0.71 |
| Nicheformer_batch | 0.79 | 0.64 | 0.57 | 0.75 | 0.73 | 0.67 | 0.74 | 0.70 |
| scGPT_batch | 0.79 | 0.61 | 0.59 | 0.66 | 0.74 | 0.66 | 0.70 | 0.68 |
| scCello_batch | 0.80 | 0.65 | 0.68 | 0.53 | 0.72 | 0.71 | 0.63 | 0.67 |
| UCE_batch | 0.76 | 0.56 | 0.58 | 0.79 | 0.54 | 0.63 | 0.66 | 0.65 |
| Geneformer_batch | 0.68 | 0.43 | 0.52 | 0.78 | 0.72 | 0.54 | 0.75 | 0.65 |
| scFoundation | 0.80 | 0.65 | 0.57 | 0.39 | 0.78 | 0.67 | 0.58 | 0.63 |
| GeneCompass_batch | 0.60 | 0.41 | 0.46 | 0.78 | 0.74 | 0.49 | 0.76 | 0.62 |
| PCA | 0.76 | 0.54 | 0.59 | 0.38 | 0.82 | 0.63 | 0.60 | 0.62 |
| CellPLM | 0.78 | 0.63 | 0.56 | 0.33 | 0.79 | 0.66 | 0.56 | 0.61 |
| scGPT | 0.79 | 0.61 | 0.54 | 0.30 | 0.80 | 0.64 | 0.55 | 0.60 |
| GeneCompass | 0.59 | 0.40 | 0.45 | 0.60 | 0.76 | 0.48 | 0.68 | 0.58 |
| Nicheformer | 0.79 | 0.63 | 0.54 | 0.28 | 0.72 | 0.65 | 0.50 | 0.58 |
| UCE | 0.76 | 0.54 | 0.57 | 0.50 | 0.41 | 0.62 | 0.46 | 0.54 |
| Geneformer | 0.68 | 0.43 | 0.49 | 0.43 | 0.65 | 0.53 | 0.54 | 0.54 |
| Scanorama | 0.74 | 0.50 | 0.50 | 0.18 | 0.75 | 0.58 | 0.46 | 0.52 |

**Figure S13. Comparison of embedding from scFMs before and after batch-centralization on the HLCA dataset with RNA source as batch source. a**, Comparison of UMAP visualizations of raw and batch-centered embeddings derived from scFMs. **b**, Detailed scIB scores of traditional methods and scFMs.

**a**

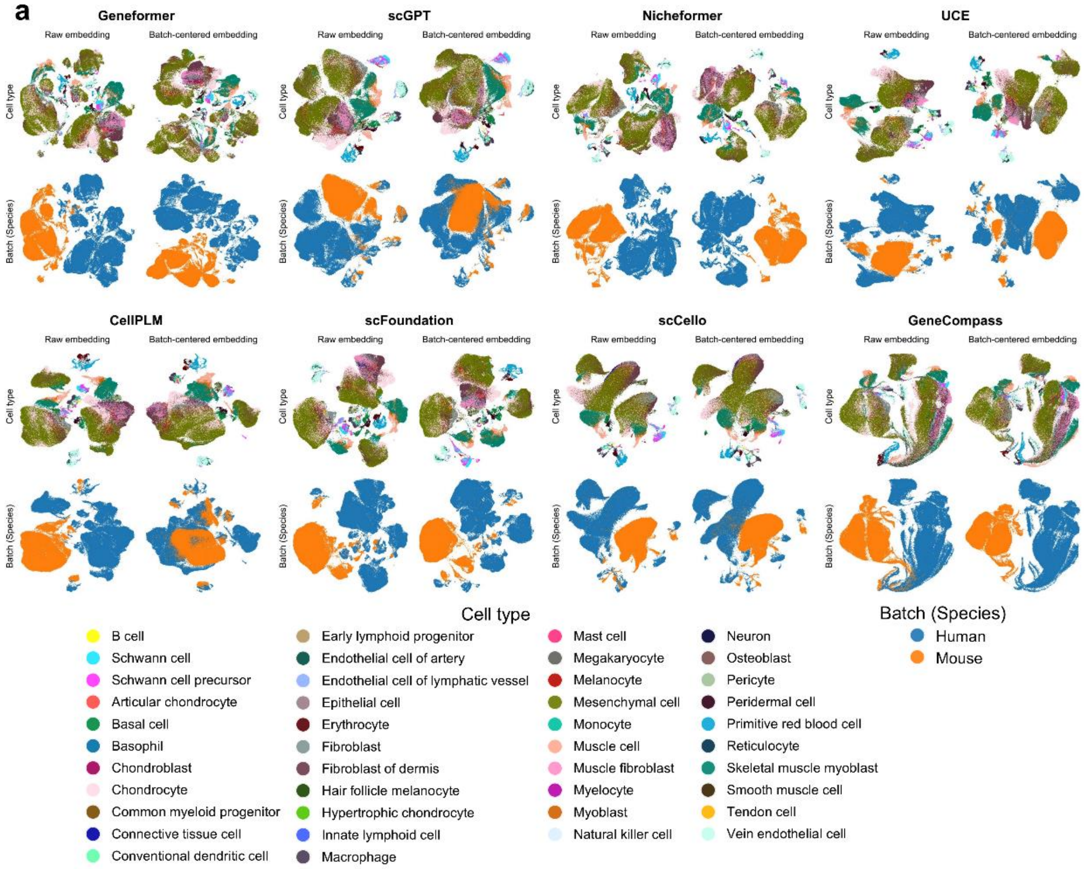


**b**

| Method | Bio conservation | | | Batch correction | | Aggregate score | | |
|---|---|---|---|---|---|---|---|---|
| | Leiden NMI | Leiden ARI | Silhouette label | BRAS | Graph connectivity | Bio conservation | Batch correction | Total |
| scVI | 0.50 | 0.19 | 0.51 | 0.77 | 0.85 | 0.40 | 0.81 | 0.60 |
| Harmony | 0.51 | 0.19 | 0.51 | 0.81 | 0.72 | 0.40 | 0.77 | 0.58 |
| scFoundation_batch | 0.43 | 0.18 | 0.49 | 0.77 | 0.79 | 0.37 | 0.78 | 0.57 |
| scGPT_batch | 0.41 | 0.18 | 0.48 | 0.79 | 0.75 | 0.36 | 0.77 | 0.56 |
| CellPLM_batch | 0.40 | 0.10 | 0.47 | 0.77 | 0.78 | 0.32 | 0.77 | 0.55 |
| UCE_batch | 0.44 | 0.20 | 0.48 | 0.80 | 0.64 | 0.37 | 0.72 | 0.55 |
| scGPT | 0.45 | 0.22 | 0.48 | 0.62 | 0.76 | 0.38 | 0.69 | 0.54 |
| scFoundation | 0.44 | 0.21 | 0.50 | 0.62 | 0.75 | 0.38 | 0.68 | 0.53 |
| UCE | 0.46 | 0.23 | 0.50 | 0.69 | 0.62 | 0.40 | 0.65 | 0.53 |
| Geneformer_batch | 0.36 | 0.11 | 0.48 | 0.79 | 0.64 | 0.32 | 0.72 | 0.52 |
| scCello_batch | 0.40 | 0.14 | 0.45 | 0.69 | 0.66 | 0.33 | 0.67 | 0.50 |
| scCello | 0.42 | 0.18 | 0.46 | 0.62 | 0.67 | 0.35 | 0.65 | 0.50 |
| PCA | 0.47 | 0.11 | 0.50 | 0.61 | 0.65 | 0.36 | 0.63 | 0.49 |
| CellPLM | 0.40 | 0.11 | 0.48 | 0.59 | 0.68 | 0.33 | 0.64 | 0.49 |
| Nicheformer_batch | 0.37 | 0.09 | 0.47 | 0.71 | 0.60 | 0.31 | 0.66 | 0.48 |
| GeneCompass_batch | 0.32 | 0.09 | 0.45 | 0.77 | 0.52 | 0.29 | 0.65 | 0.47 |
| Geneformer | 0.35 | 0.06 | 0.47 | 0.63 | 0.59 | 0.29 | 0.61 | 0.45 |
| GeneCompass | 0.31 | 0.08 | 0.44 | 0.58 | 0.53 | 0.28 | 0.56 | 0.42 |
| Scanorama | 0.44 | 0.09 | 0.45 | 0.40 | 0.60 | 0.32 | 0.50 | 0.41 |
| Nicheformer | 0.38 | 0.11 | 0.45 | 0.43 | 0.59 | 0.31 | 0.51 | 0.41 |

**Figure S14. Comparison of embedding from scFMs before and after batch-centralization on the limb dataset. a**, Comparison of UMAP visualizations of raw and batch-centered embeddings derived from scFMs. **b**, Detailed scIB scores of traditional methods and scFMs.

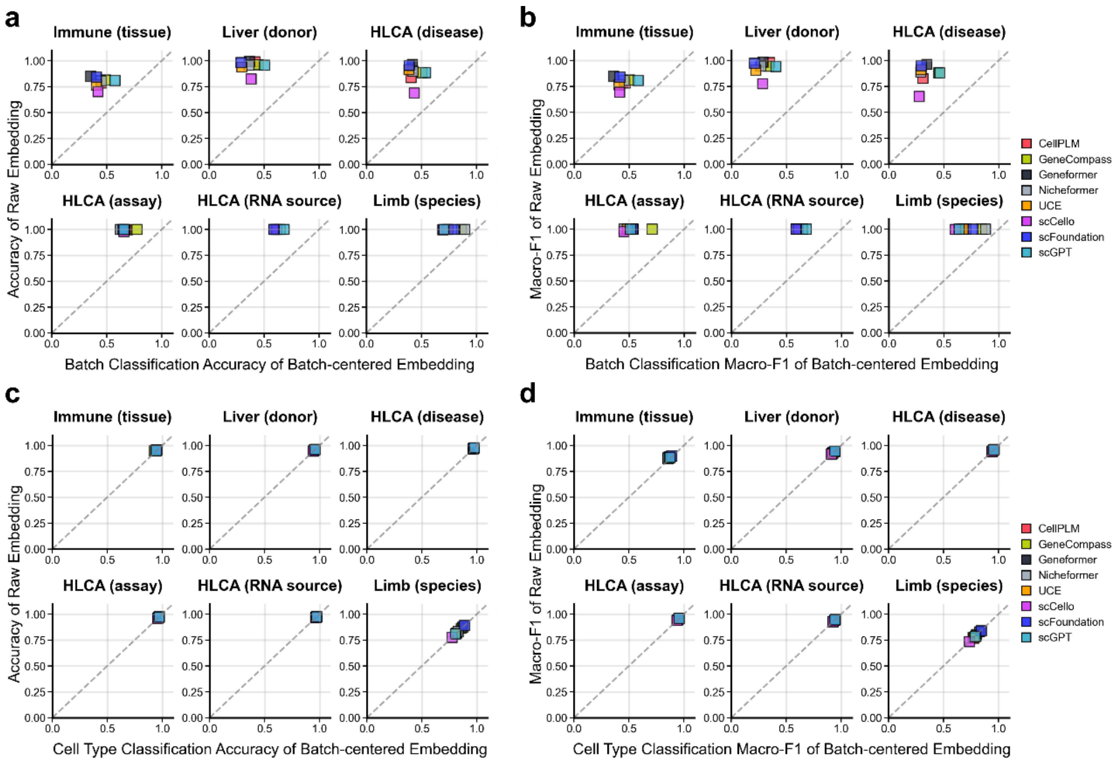


**Figure S15. Comparison of probing analysis results before and after batch-centralization. a** and **b**, Changes in batch classification accuracy (**a**) and macro-F1 score (**b**) before and after batch-centralization across six datasets for eight scFMs. **c** and **d**, Changes in cell-type classification accuracy (**c**) and macro-F1 score (**d**) before and after batch-centralization across six datasets for eight scFMs.

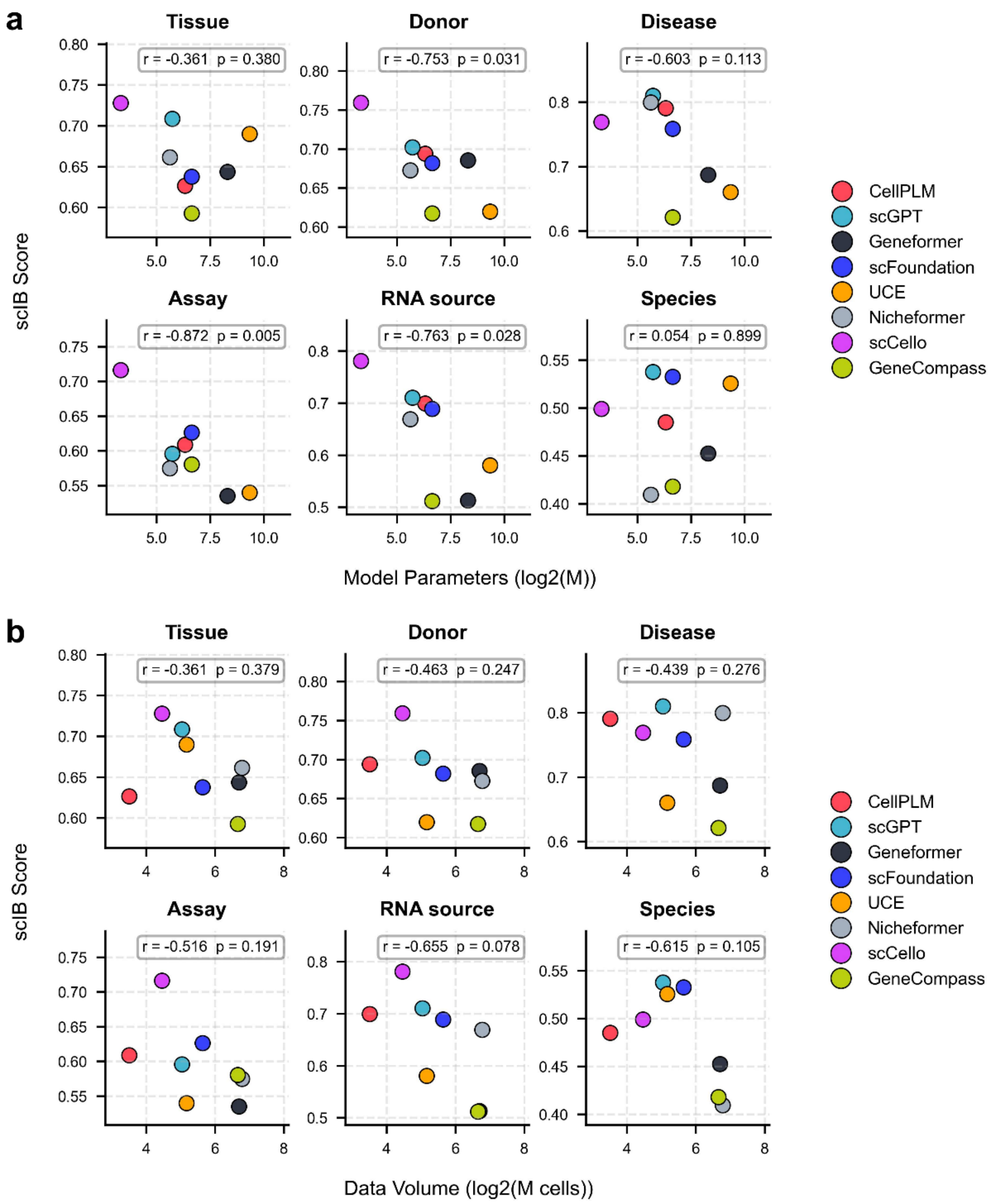


**Figure S16. Evaluation of scaling laws of current scFMs. a** and **b**, Comparison of scIB score with model parameters (**a**) or data volume (**b**) of scFMs. The Pearson correlation coefficient and *P*-value of each dataset are calculated.

**Supplementary Table S1. Summary of all evaluation datasets and their key characteristics.**

| Data name | Batch source | Cell number | Data link | Reference |
|---|---|---|---|---|
| Liver (donor) | 16 donors | 68429 | https://cellxgene.cziscience.com/collections/ff69f0ee-fef6-4895-9f48-6c64a68c8289 | 10.1097/HC9.0000000000000813 |
| Immune (tissue) | 10 tissues | 85588 | https://cellxgene.cziscience.com/collections/cc431242-35ea-41e1-a100-41e0dec2665b | 10.1038/s41590-025-02241-4 |
| HLCA (disease) | 4 disease and normal | 112818 | https://cellxgene.cziscience.com/collections/6f6d381a-7701-4781-935c-db10d30de293 | 10.1038/s41591-023-02327-2 |
| HLCA (assay) | 2 assay:<br>10x 3' v2 and Drop-seq | 54414 | | |
| HLCA (RNA source) | 2 source:<br>single-cell and single-nucles | 73915 | | |
| Limb (species) | 2 species:<br>human and mouse | 200866 | https://cellxgene.cziscience.com/collections/4fefa187-5d14-4f1e-915b-c892ed320aab | 10.1038/s41586-023-06806-x |

**Supplementary Table S2. Summary of the basic information of current scFMs.** MLM, masked language modeling; KL, Kullback-Leibler divergence; SCL, Supervised contrastive loss.

| Model | Time | Params | Data size (cell) | Tokenizer strategy | Input embedding | Architecture | Loss | Batch correction |
|---|---|---|---|---|---|---|---|---|
| scBERT | 22-09 | 8.4M | 1.1M | Binning | expression + geneID | Encoder (BERT) | MLM | - |
| Geneformer | 23-05 | 13.5M | 27.4M | Ordering | geneID + position | Encoder (BERT) | MLM | - |
| scGPT | 23-05 | 53M | 33M | Binning | expression + geneID | Encoder | MLM | - |
| scFoundation | 23-06 | 100M | 50M | Value projection | expression + geneID | Encoder-decoder | MLM | - |
| CellPLM | 23-10 | 80M | 11.4M | Value projection | cell + position | Encoder-decoder | MLM + KL | Batch embedding in decoder |
| UCE | 23-11 | 650M | 36M | Ordering | geneID + position | Encoder (BERT) | MLM | - |
| Nicheformer | 24-04 | 49.3M | 110M | Ordering | geneID + position | Encoder (BERT) | MLM | Modality/Organism/Assay token |
| scCello | 24-08 | 10M | 22M | Same as Geneformer | | | MLM + SCL | Contrastive learning using cell type |
| Geneformer2 | 25-06 | 316M | 104M | Ordering | geneID + position | Encoder (BERT) | MLM | - |
| CellFM | 25-05 | 800M | 100M | Value projection | expression + geneID | Encoder (BERT) | MLM | - |
| TEDDY | 25-03 | 400M | 116M | Same as Geneformer and scGPT | | | MLM | - |
| Nephrobase Cell+ | 25-10 | 1B | 39.5M | Value projection | expression + geneID | Encoder-decoder | Combined loss | Contrastive learning using cell type; Adversarial learning |
